# Fitting quantum machine learning potentials to experimental free energy data: Predicting tautomer ratios in solution

**DOI:** 10.1101/2020.10.24.353318

**Authors:** Marcus Wieder, Josh Fass, John D. Chodera

**Author notes:** contributed equally to this work (Marcus Wieder, Josh Fass). Relay Therapeutics, Cambridge, MA 02139, USA. **For correspondence:** (MW).

## Abstract

The computation of tautomer ratios of druglike molecules is enormously important in computer-aided drug discovery, as over a quarter of all approved drugs can populate multiple tautomeric species in solution. Unfortunately, accurate calculations of aqueous tautomer ratios—the degree to which these species must be penalized in order to correctly account for tautomers in modeling binding for computer-aided drug discovery—is surprisingly difficult. While quantum chemical approaches to computing aqueous tautomer ratios using continuum solvent models and rigid-rotor harmonic-oscillator thermochemistry are currently state of the art, these methods are still surprisingly inaccurate despite their enormous computational expense. Here, we show that a major source of this inaccuracy lies in the breakdown of the standard approach to accounting for quantum chemical thermochemistry using rigid rotor harmonic oscillator (RRHO) approximations, which are frustrated by the complex conformational landscape introduced by the migration of double bonds, creation of stereocenters, and introduction of multiple conformations separated by low energetic barriers induced by migration of a single proton. Using quantum machine learning (QML) methods that allow us to compute potential energies with quantum chemical accuracy at a fraction of the cost, we show how rigorous relative alchemical free energy calculations can be used to compute tautomer ratios in vacuum free from the limitations introduced by RRHO approximations. Furthermore, since the parameters of QML methods are tunable, we show how we can train these models to correct limitations in the underlying learned quantum chemical potential energy surface using free energies, enabling these methods to learn to generalize tautomer free energies across a broader range of predictions.

## Introduction

The most common form of tautomerism, *prototropic tautomerism* describes the reversible structural isomerism involving the sequential processes of bond cleavage, skeletal bond migration and bond reformation in which a H^+^ is transferred [1]. Numerous chemical groups can show prototropic tautomerism. Common examples include keto-enol (shown in Figure 2), amide/imidic acid, lactam/lactim, and amine/imine tautomerism [2].

### Tautomerism influences many aspects of chemistry and biology

Tautomerism adds a level of mutability to the static picture of chemical compounds. The widespread usage of notating and representing chemical structure as undirected graphs with atoms as nodes and bonds as edges (and bond types annotated as edge attributes like single, double and triple bond) dominates the field of chemistry (with the notable exception of quantum chemistry) [3]. Tautomers make a satisfying description of chemical identity difficult in such a framework, since they can instantaneously change their double bond pattern and connectivity table if confronted with simple changes in environment conditions or exist as multiple structures with different connection table and bond pattern at the same time [4].

The change in the chemical structure between different tautomeric forms of a molecule is accompanied by changes in physico-chemical properties. By virtue of the movement of a single proton and the rearrangement of double bonds, tautomerism can significantly alter a molecule’s polarity, hydrogen bonding pattern, its role in nucleophilic/electrophilic reactions, and a wide variety of physical properties such as partition coefficients, solubilities, and pKa [5, 6].

Tautomerism can also alter molecular recognition, making it an important consideration for supramolecular chemistry. To optimizing hydrogen bond patterns between a ligand and a binding site tautomerismn has to be considered. In a theoretical study of all synthetic, oral drugs approved and/or marketed since 1937, it has been found that 26% exist as an average of three tautomers [7]. While tautomerism remains an important phenomena in organic chemistry, it has not gained much appreciation in other scientific fields.

The typical small free energy difference between tautomers poses additional challenges for proteinligand recognition. Local charged or polar groups in the protein binding pocket can shift the tautomer ratio and result in dominant tautomers in complex otherwise not present in solution [3]. For this reason the elucidation of the *dominant* tautomer in each environment is not enough — without the knowledge about the ratio (i.e. the free energy difference) between tautomeric forms in the corresponding phase a correct description of the experimental (i.e. macroscopic) binding affinity might not be possible. To illustrate this, one might think about two extreme cases: one in which the tautomeric free energy difference in solution is 10 kcal/mol and one in which it is 1.0 kcal/mol. It seems unlikely that the free energy difference of 10 kcal/mol will be compensated by the protein binding event (therefore one could ignore the unlikely *other* tautomer form), but a tautomeric free energy difference of 1 kcal/mol could easily be compensated. Examples for this effect — changing tautomer ratios between environments — are numerous for vacuum and solvent phase. One example is the neutral 2-hydroxypyridine(2-HPY)/2-pyridone(2-PY) tautomer. The gas phase internal energy difference between the two tautomers is ∼0.7 kcal/mol in favor of the enol form (2-HPY), while in water an internal energy difference of 2.8 kcal/mol was reported in favour of 2-PY. In cyclohexane, both tautomer coexist in comparable concentration while the 2-PY is dominant in solid state and polar solvents [8].

The example above assumed two tautomeric forms — if there were multiple tautomeric forms, each with small free energy differences, using a single dominant tautomer in solvent and complex may still represent only a minor fraction of the true equilibrium tautomer distribution.

### The study of tautomer ratios requires sophisticated experimental and computational approaches

The experimental study of tautomer ratios is highly challenging [9]. The small free energy difference and low reaction barrier between tautomers—as well as their short interconversion time—can make it nearly impossible to study specific tautomers in isolation. Recently, efforts have been made to collect some of the experimental data and curate these in publicly available databases, specifically Wahl and Sander [10] and Dhaked et al. [11].

In the absence of a reliable, cheap, and fast experimental protocol to characterize tautomeric ratios for molecules, there is a need for computational methods to fill the gap. And — even if such a method exists — predicting tautomer ratios of molecules that are not yet synthesized motivates the development of theoretical models. Computational methods themselves have a great need for defined tautomer ratios. Most computational methods use data structures in which bond types and/or hydrogen positions have to be assigned *a priori* and remain static during the calculation, introducing significant errors if an incorrect dominant tautomer is chosen [12].

The third round of the Statistical Assessment of the Modeling of Proteins and Ligands (SAMPL2) challenge included the blind prediction of tautomer ratios and provided an interesting comparison between different computational methods [13]. Most of the 20 submissions were using implicit solvent models and *ab initio* or DFT methods in combination with a thermodynamic cycle (as shown in Figure 1) to evaluate tautomeric free energy difference [13]. However, as stated in Martin [14], “In summary, although quantum chemical calculations provide much insight into the relative energies of tautomers, there appears to be no consensus on the optimal method”. For the 20 tautomer pairs investigated in the SAMPL2 challenge (this includes 13 tautomer pairs for which tautomer ratios were provided beforehand) the three best performing methods reported an RMSE of 2.0 [15], 2.5 [16] and 2.9 kcal/mol [17]. While these results are impressive it also shows that there is clearly room for improvement. And, maybe even more importantly, the SAMPL2 challenge showed the need for investigating a wider variety of tautomer transitions to draw general conclusions about best practices and methods for tautomer predictions [13]. The excellent review of Nagy concluded that tautomer relative free energies are sensitive to “the applied level of theory, the basis set used both in geometry optimization and […] single point calculations, consideration of the thermal corrections […] and the way of calculating the relative solvation free energy” [18]. We’d like to add “quality of 3D coordinates” to these issues and discuss them in the following.

### Standard approaches to quantum chemical calculations of tautomer ratios introduce significant errors

The tautomeric free energy difference in solution 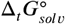 can be calculated from the standard-state Gibbs free energy in aqueous phase 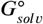 of the product and educt of the corresponding tautomer reaction, which itself is calculated as the sum of the gas-phase standard-state free energy 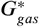 and the standard-state transfer free energy 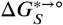, shown as a thermodynamic cycle in Figure 1.

**Figure 1.**
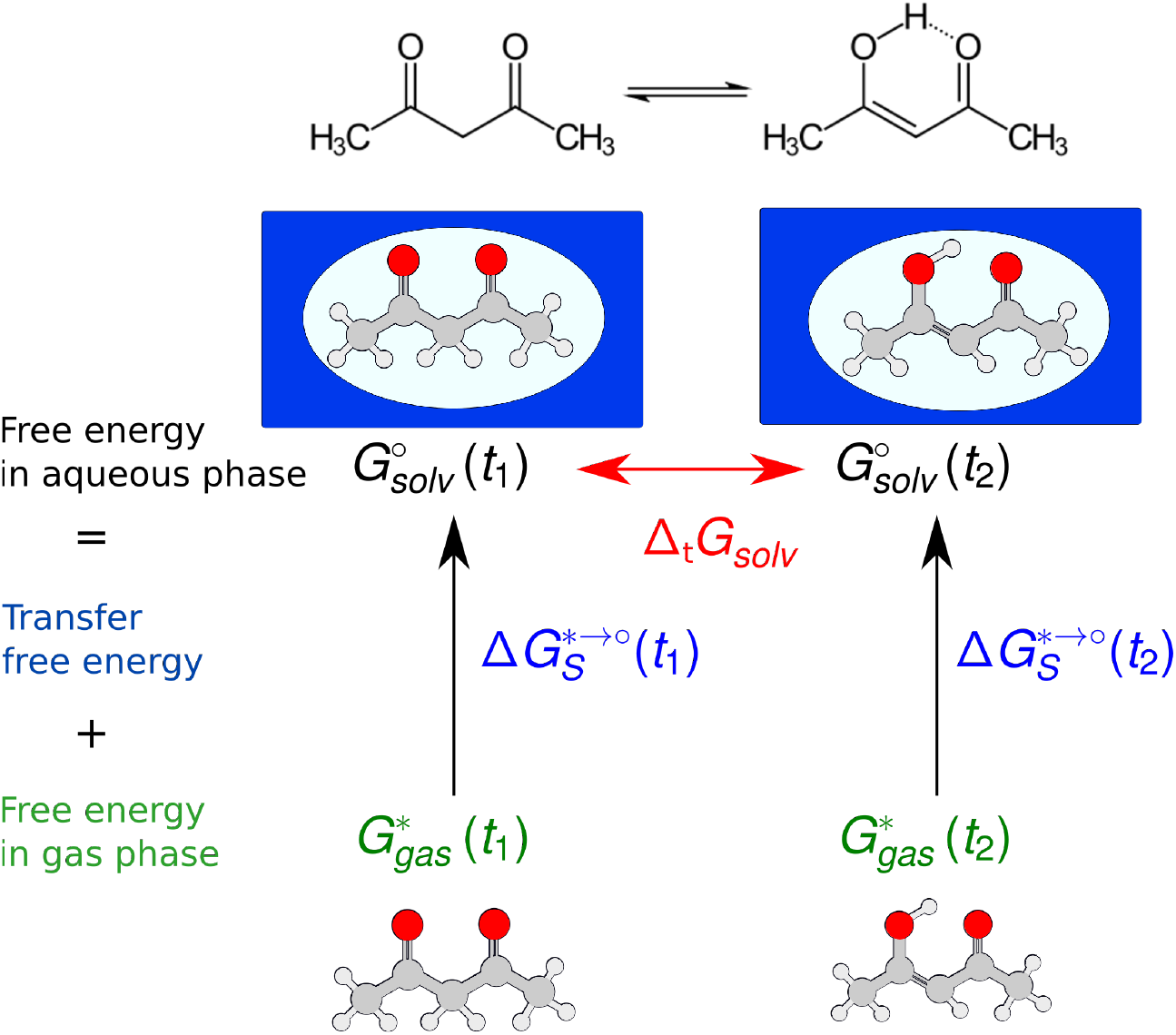
Tautomeric free energy differences in solution are typically calculated using a thermodynamic cycle, independent of the level of theory that is used to obtain the individual terms. A typical quantum chemistry protocol calculates the free energy in aqueous phase 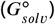 as the sum of the free energy in gas phase (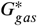; calculated using the ideal gas RRHO approximation) and the standard state transfer free free energy (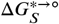; obtained using a continuum solvation model). The *tautomeric free energy difference in solution* 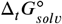 is then calculated as the difference between 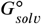 for two tautomers.

**Figure 2.**
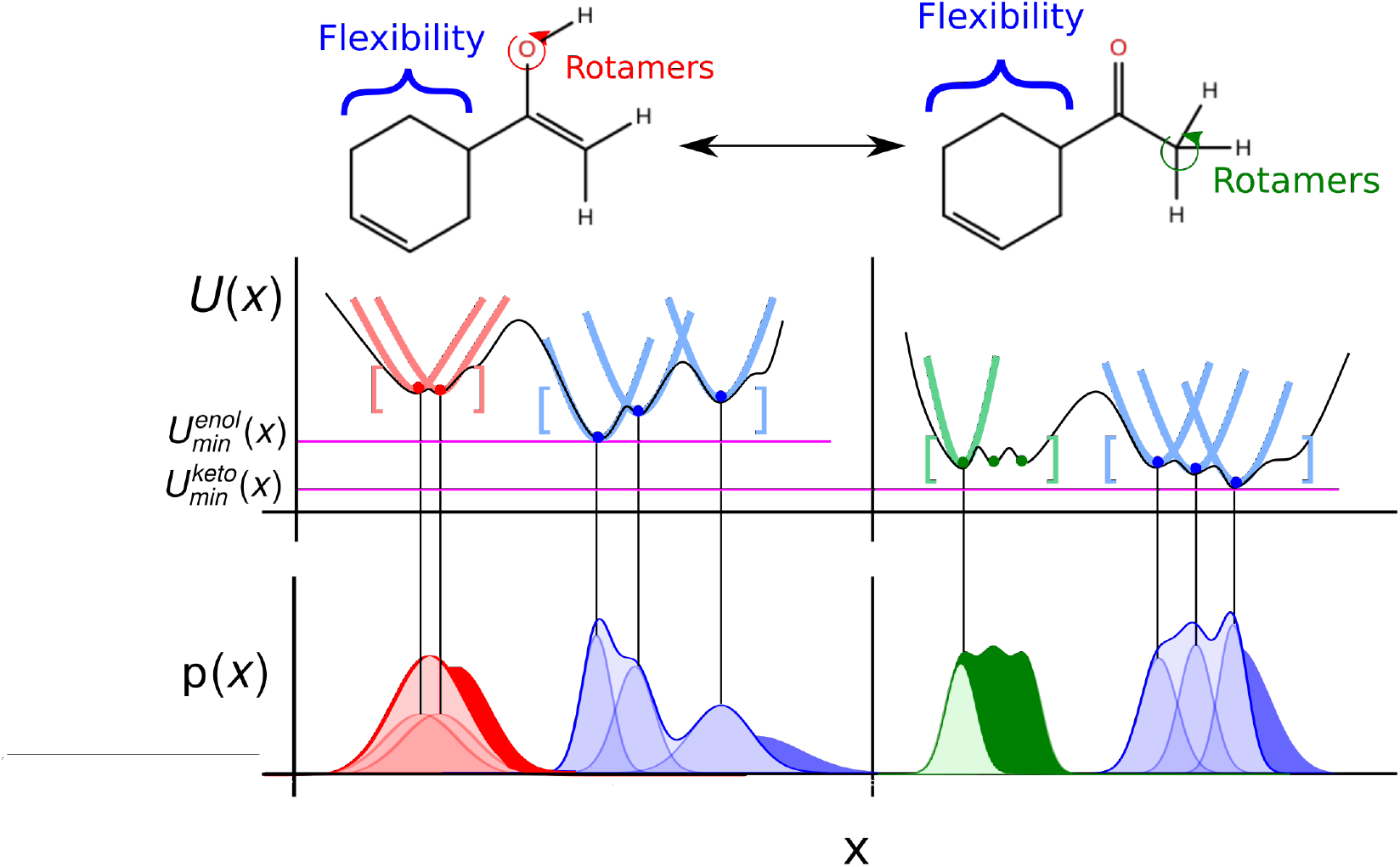
The rigid-rotor harmonic oscillator (RRHO) approach constructs a partition function using the curvature of the potential energy landscape at a minimum modelling all bonded terms as harmonic. The enol and keto form of a molecules (in the enol form: 1-(cyclohex-3-en-1-yl)ethen-1-ol) is shown and the main conformational degrees of freedom highlighted. The middle panel shows how the use of a local and harmonic approximation to the partition function can approximate the overall potential energy landscape, if all relevant minimum conformations are enumerated. The lower panel shows the probability density resulting from the harmonic approximations, with solid colored regions the difference between the true and approximated probability density resulting from anharmonicity and/or bonded terms that would better be modeled using a hindered/free rotor.

**Figure 3.**
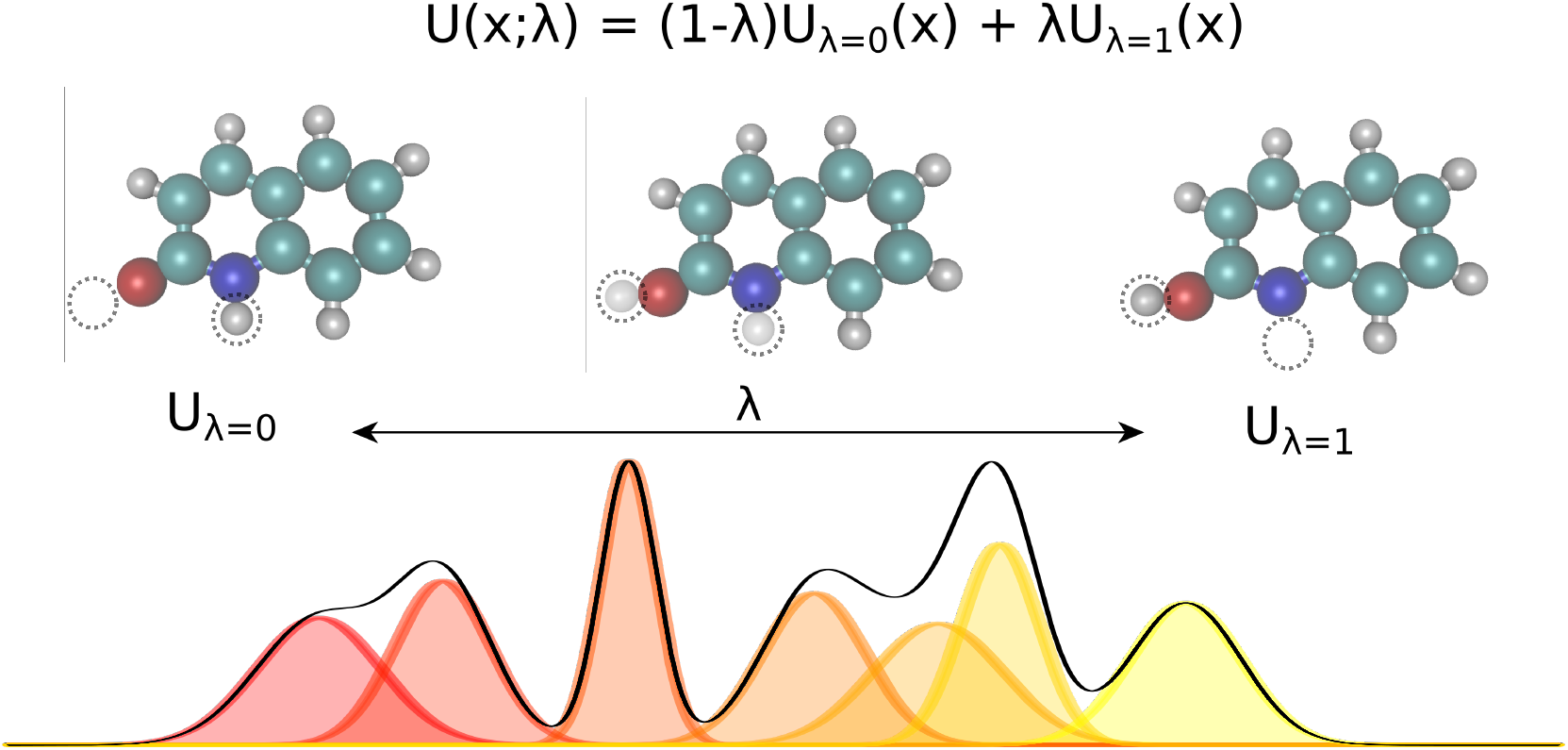
Alchemical relative free energy calculations with quantum machine learning (QML) potentials like ANI can rigorously compute free energy differences. Linearly interpolating between two potential energy functions using the alchemical parameter *Jc* enables the sampling of bridging distributions between the physical endstates representing the two tautomers. Here, noninteracting dummy atoms—indicated by the empty circle at each endstate—are included to facilitate this transformation. In this work, we present the application of this concept to calculate relative free energies of tautomer pairs in vacuum.

The standard-state Gibbs free energy is calculated using a quantum chemistry estimate for the electronic energy and, based on its potential energy surface, a statistical mechanics approximation to estimate its thermodynamic properties. While the accuracy of the electronic energy and the transfer free energy Δ*G*_*s*_ is dependent on the chosen method and can introduce significant errors in the following we want to concentrate on the approximations made by the thermochemical correction to obtain the gas phase free energy: the rigid rotor harmonic oscillator (RRHO) approximation and the use of single or multiple minimum conformations to generate a discrete partition function. In the following the standard-state of all the thermodynamic properties should be implied and will not be added to the notation.

A commonly used approach for the zero point energy (ZPE) and thermal contributions is based on the ideal gas RRHO, assuming that the molecule is basically rigid and its internal motions comprise only vibrations of small amplitude where the potential energy surface can be approximated as harmonic around a local energy minimum. This assumption leads to an analytical approximation that describes the local configuration space around a single minimum based on the local curvature of the potential energy surface (Figure 2) [19].

Errors are introduced because the harmonic oscillator approximation assumes that, for each normal mode, the potential energy associated with the molecule’s distortion from the equilibrium geometry has a harmonic form. Especially low-frequency torsional modes would be more appropriately treated as hindered internal rotations at higher temperatures. This can lead to a significant underestimate of the configurational entropy, ignoring the contributions of multiple energy wells [20]. The correct treatment of such low-frequency modes in the analytical rotational entropy part of the RRHO partition function can add considerable computational cost, since the numerical solution requires the calculation of the full torsion potential (periodicity and barrier heights) [21].

Another, related shortcoming of the RRHO approach is the focus on a single minimum conformation. Assuming that the correct global minimum has been identified this approach ignores the configurational entropy of all other conformations to the partition function. Methods like the ‘mining minima’ approach can help to mitigate this problem and construct a partition function using local configurational integrals from multiple minimum energy wells [20]. The standard state free energy *G*^°^ of a molecule can then be calculated for each of the minimum conformations separately and combined as follows

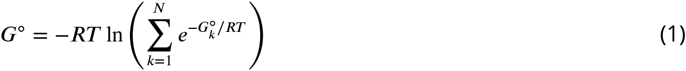

to obtain a weighted free energy [22, 23]. The free energy obtained in Equation 1 mitigates the shortcomings of the RRHO approach to model torsions with periodicity of 1 since — as long as the different minima of the torsion are generated as conformations — these are included in the sum over all conformations. But the configurational integral is still only taken over the minimum conformation and the contribution of its energy well, ignoring anharmonicity in the bonded terms and contributions from conformations outside the minimum energy wells.

### Alchemical relative free energy calculations with machine learning potentials can compute true tautomeric free energy differences, including all classical statistical mechanical effects

Limitations of the ideal gas RRHO approximation to the free energy and ZPE correction, challenges in enumeration of local minimum conformations, and the consistent treatment of internal/external rotational symmetry, as well as approximations in the continuum solvent model can lead to errors in the free energy that are difficult to detect and correct. The use of molecular dynamics simulations, explicit solvent molecules, and a rigorous classical treatment of nuclear motion to sample independent conformations from an equilibrium distribution can overcome the above mentioned challenges. To describe tautomer free energies in solution this would require computationally prohibitive QM/MM calculations in which the molecule of interest is treated quantum mechanically and the solute treated with molecular mechanics.

One of the most exciting developments in recent years has been the introduction of fast, efficient, accurate and transferable quantum machine learning (QML) potentials (e.g. ANI [24], PhysNet [25], and SchNet [26]) to model organic molecules. QML potentials can be used to compute QM-based Hamiltonians and—given sufficient and appropriate training data—are able to reproduce electronic energies and forces without loss of accuracy but with orders of magnitude less computational cost than the QM methods they aim to reproduce. QML potentials have been successfully applied to molecular dynamics (MD) and Monte Carlo (MC) simulations [27]. Here, we present the application of a machine learning potential for the calculation of *alchemical free energies* for tautomer pairs in the gas phase.

We begin by investigating the accuracy of a current, state of the art approach to calculate tautomer ratios on a set of 468 tautomer pairs selected from the publicly available *Tautobase* dataset [10] spanning different tautomer reactions, number of atoms and functional groups. We are using a popular DFT functional (B3LYP) and basis sets [aug-cc-pVTZ and 6-31G(d)] [28–32] for the calculation of the electronic energy and a continuum solvation model (SMD [33]) to model the transfer free energy.

Furthermore we are investigating the effect of a more rigorous statistical mechanics treatment of the gas phase free energies. To investigate this effect we calculate tautomeric free energy difference in vacuum 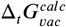 using (1) alchemical relative free energy calculations and (2) multiple minimum conformations in combination with the RRHO approximation (as used in the quantum chemistry approach). To enable a direct comparison between the two approaches we are using a QML potential in both calculations.

Since we are interested in tautomeric free energy differences in *solution* we investigate the possibility to optimize the QML parmaters to include crucial solvent effects. The framework we have developed to perform alchemical relative free energy calculations enables us to obtain a relative free energy estimate that can be optimized with respect to the QML parameters. We are using a small training set of experimentally obtained tautomer free energies in *solution* 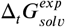 and importance sampling to obtain reweighted 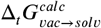 with the optimized parameters.

Here we report the first large scale investigation of tautomer ratios using quantum chemical calculations and machine learning potentials in combination with alchemical relative free energy calculations spanning a large chemical space and 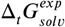 values.

### Standard quantum chemistry methods predict tautomer ratios with an RMSE of kcal/mol for a large set of tautomer pairs

Calculating tautomeric free energy differences in solution 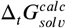 is a challenging task since its success depends on a highly accurate estimate of both the intrinsic free energy difference between tautomer pairs as well as the transfer free energy.

We calculated the tautomeric free energy difference 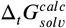 for 468 tautomer pairs (460 if the basis set 6-31G(d) was used) and compared the results with the experimentally obtained tautomer ratios in solution (expressed as free energy difference in solution 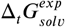) in Figure 4.

**Figure 4.**
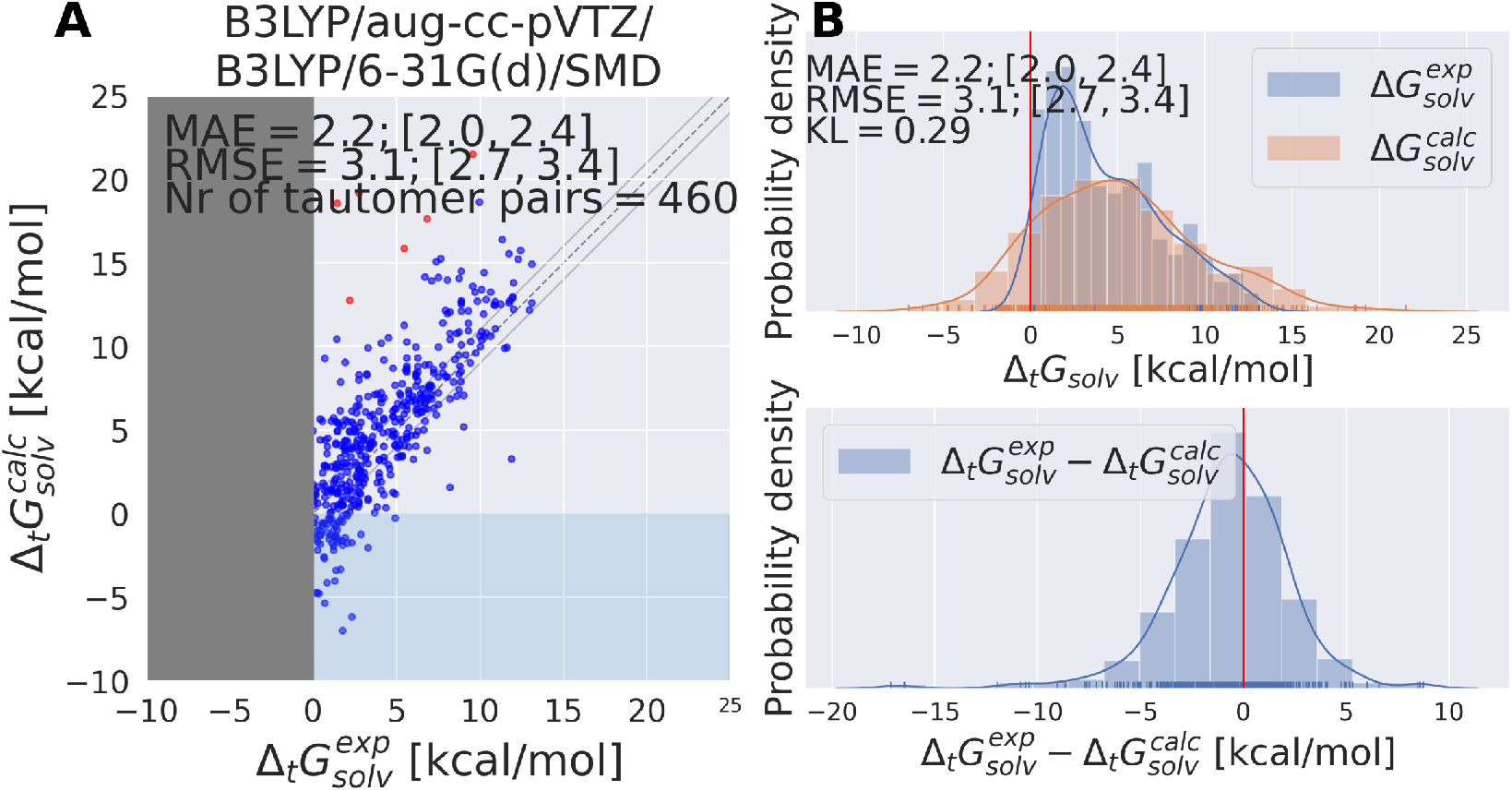
State of the art quantum chemistry calculations are able to calculate tautomeric free energy differences 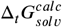 with a RMSE of 3.1 kcal/mol. The direction of the tautomer reaction is chosen so that the experimentally obtained tautomeric free energy difference 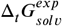 is always positive. Panel **A** shows 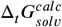 as the difference between the sum of the gas phase free energy and transfer free energy for each tautomer pair plotted against the experimental tautomeric free energy difference in solution 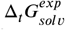. B3LYP/aug-cc-pVTZ is used for the gas phase geometry optimization and single point energy calculation, the ideal gas RRHO approximation is used to calculate the thermal corrections. The transfer free energy is calculated on B3LYP/aug-cc-pVTZ optimized geometries in their respective phase using B3LYP/6-31G(d) and SMD. Values in quadrant II indicate calculations that assigned the wrong dominant tautomer species (different sign of 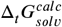 and 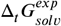). Red dots indicate tautomer pairs with more than 10 kcal/mol absolute error between 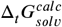 and 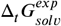. These are separately shown in Table S.I.1. In Panel **B**, the top panel shows the kernel density estimate (KDE) and histogram of 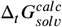 and 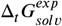. The red line indicates zero free energy difference (equipopulated free energies). In the lower panel the KDE of the difference between 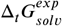 and 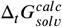 is shown. MAE and RMSE are reported in units of kcal/mol. Quantities in brackets [X;Y] denote 95% confidence intervals. The Kullback-Leibler divergence (KL) was calculated using 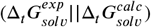.

Three quantum chemistry approaches were used to calculate these values:

- **B3LYP/aug-cc-pVTZ/B3LYP/aug-cc-pVTZ/SMD:** Multiple conformations were generated for each tautomer, geometry optimization and energy calculations were performed with B3LYP/aug-cc-pVTZ in gas phase and solution (using the SMD continuum solvation model) and 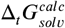 was calculated as the difference between the free energies in aqueous phase for the individual tautomers. This approach will be abbreviated as B3LYP/aug-cc-pVTZ/B3LYP/aug-cc-pVTZ/SMD in the following. The RMSE between 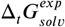 and 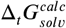 for the dataset is 3.4 [3.0;3.7] kcal/mol (the quantities [X;Y] denote a 95% confidence interval).
- **B3LYP/aug-cc-pVTZ/B3LYP/6-31G(d)/SMD** Multiple conformations were generated for each tautomer, conformations were optimizing with B3LYP/aug-cc-pVTZ in gas phase and solution (using the SMD continuum solvation model). Transfer free energy was calculated using B3LYP/6-31G(d)/SMD. 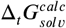 was calculated as the difference between the free energy in aqueous phase for the individual tautomers and conformations. The RMSE between 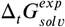 and 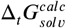 for the QM dataset is 3.1 [2.7;3.4] kcal/mol. This approach will be subsequently called B3LYP/aug-cc-pVTZ/B3LYP/6-31G(d)/SMD
- **B3LYP/aug-cc-pVTZ/SMD** Generating multiple conformations, optimizing with B3LYP/aug-cc-pVTZ in solution phase (using the SMD solvation model) and calculating relative solvation free energy 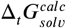 directly as the difference between the free energy in aqueous phase (on the solution phase geometry). In this case, the free energy in aqueous phase is *not* obtained through a thermodynamic cycle, but the frequency calculation and thermochemistry calculations are performed with the continuum solvation model [34]. The individual free energy in aqueous phase 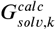 for conformation *k* are averaged to obtain the final 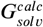. The RMSE between 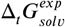 and 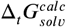 is 3.3 [3.0;3.7] kcal/mol. This approach will be subsequently called B3LYP/aug-cc-pVTZ/SMD.

The protocol used to obtain the results described above did not explicitly account for changes in internal rotors (only changes in the point group were considered). The results including changes in internal rotors are shown in Figure S.I.3. Including internal symmetry numbers did not improve the results.

The following discussion will concentrate on the results obtained with the best performing method (B3LYP/aug-cc-pVTZ/B3LYP/6-31G(d)/SMD). The results for all other methods are shown in the Supplementary Material Section in Figure S.I.2.

The tautomeric free energy difference in solution 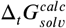 calculated with B3LYP/aug-cc-pVTZ/B3LYP/6-31G(d)/SMD are shown in Figure 4. The RMSE for the calculated values is 3.1 [2.7;3.4] kcal/mol. Overall, the method tends to overestimate the tautomeric free energy difference in solution, as can be seen from Figure 4B (lower panel). For 59 tautomer pairs (13% of the dataset) the method was not able to correctly determine the dominant tautomer species. For these 59 tautomer pairs the mean absolute error (MAE) between predicted and experimental value was 2.9 kcal/mol.

5 out of the 6 tautomer pairs with highest absolute error (above 10 kcal/mol) between experimental and calculated free energy difference have common scaffolds (values and molecules shown in the Supplementary Material Table S.I.1). tp_1668, 1669, 1670 are all based on 5-iminopyrrolidin-2-one with either nitrogen, oxygen or sulfur on position 4 of the ring. tp_1559 and tp_853 both have methoxyethylpiperidine as a common substructure.

For the three analogues based on 5-iminopyrrolidin-2-one, we note that the experimental values are estimated results based on a three-way tautomeric reaction [35]. While this might point to the unreliability of 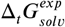, we can not draw any definitive conclusions here without repeating the underlying experiment.

### The best performing quantum chemistry method shows very good retrospective performance on the SAMPL2 tautomer set

Some of the tautomer pairs deposited in the Tautobase dataset were also part of the SAMPL2 challenge, specifically 6 out of 8 tautomer pairs of the obscure dataset (using the notation of the SAMPL2 challenge: 1A_1B, 2A_2B, 3A_3B, 4A_4B, 5A_5B, 6A_6B) and 8 out of 12 tautomer pairs of the explanatory dataset (7A_7B, 10B_10C, 10D_10C, 12D_12C, 14D_14C, 15A_15B, 15A_15C, 15B_15C) [13]. Comparing these molecules with the results of the SAMPL2 challenge—specifically with the results of the 4 participants with the best overall performance ([17], [36], [16], and [15])—helps assess the quality of the used approach. Since all four mentioned participants employed different methods, we will briefly describe the best performing methods of these publications for which results are shown in Table 1. The references to the methods used below are not cited explicitly, these can be found in the publications cited at the beginning of the following paragraphs.

**Table 1.**
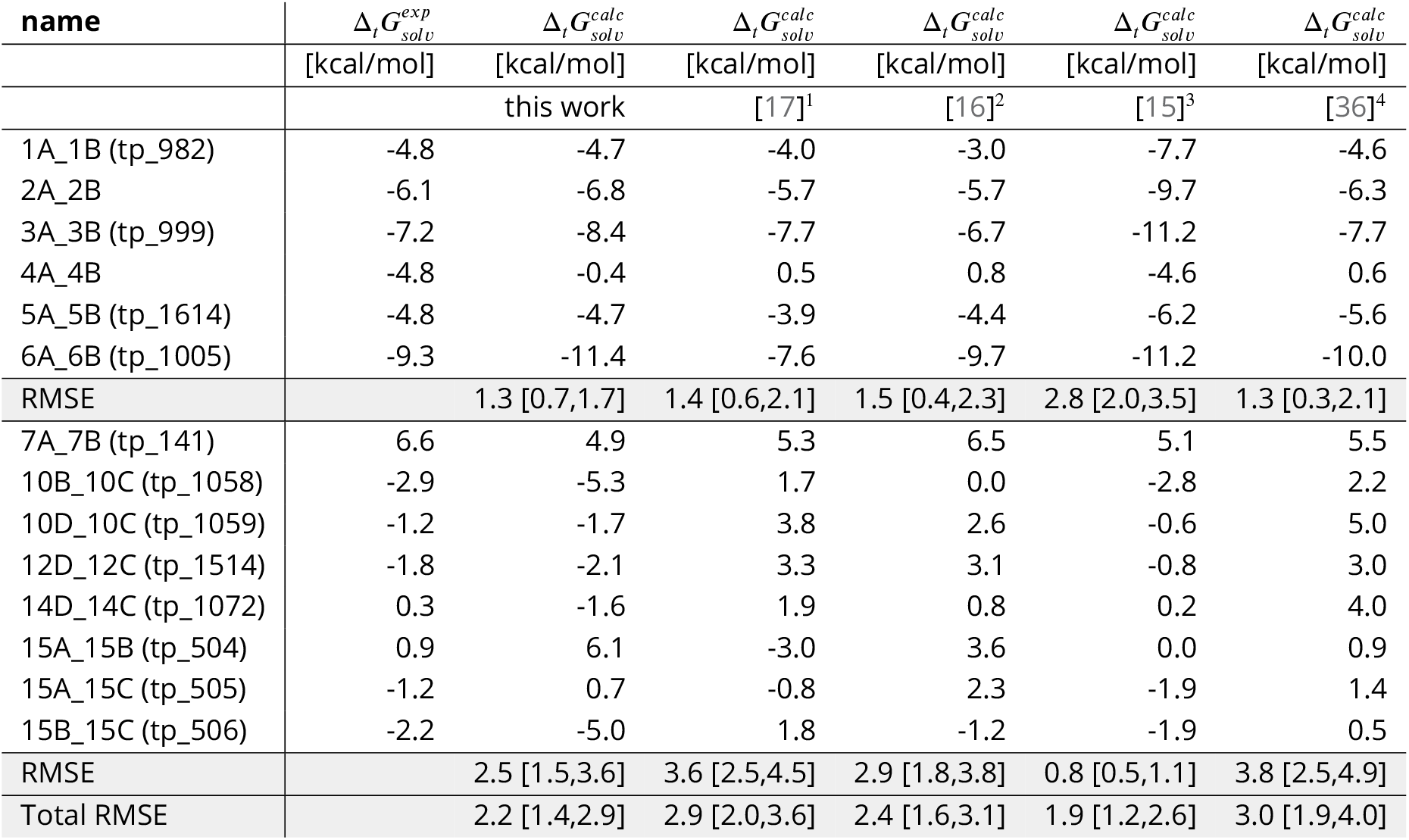
Comparison between the results of this work and the SAMPL2 challenge. All values are given in kcal/mol. The first part of the names in the ‘name’ column refers to the tautomer pair naming convention from the SAMPL2 challenge and in brackets is the name as used in this work. The calculations for Klamt and Diedenhofen [17] were performed with MP2+vib-CT-BP-TZVP, for Ribeiro et al. [16] with M06-2X/MG3S/M06-2X/6-31G(d)/SM8AD, Kast et al. [15] with MP2/aug-cc-pVDZ/EC-RISM/PSE-3, and Soteras et al. [36] with MP2/CBS+[CCSD-MP2/6-31+G(d)](d)/IEF-MST/HF/6-31G). On this subset the presented approach was the second best method based on the total RMSE. Bracketed quantities [X,Y] denote 95% confidence intervals.

Klamt and Diedenhofen [17] used BP86/TZVP DFT geometry optimization in vacuum and the COSMO solvation model. Free energy in solution was calculated from COSMO-BP86/TZVP solvation energies and MP2/QZVPP gas-phase energies. Thermal corrections (including ZPE) were obtained using BP86/TZVP ZPE.

Ribeiro et al. [16] calculated the free energy in solution as the sum of the gas-phase free energy and the transfer free energy. The gas phase free energy was calculated using M06-2X/MG3S level of theory and the molecular geometries optimized with the same method. The corresponding transfer free energy were computed at the M06-2X/6-31G(d) level of theory with the M06-2X/MG3S gas-phase geometries using the SM8, SM8AD, and SMD continuum solvation models (Table 1 shows only the results with SM8AD, which performed best).

Kast et al. [15] optimized geometries in gas and solution phase (using the polarizable continuum solvation model PCM) using B3LYP/6-311++G(d,p). Energies were calculated with EC-RISM-MP2/aug-cc-pVDZ on the optimized geometries in the corresponding phase. The Lennard-Jones parameters of the general Amber force field (GAFF) were used.

Soteras et al. [36] used the IEF-MST solvation model parameterized for HF/6-31G(G) to obtain transfer free energy values. Gas phase free energy differences were obtained by MP2 basis set extrapolation using the aug-cc-pVTZ basis set at MP2/6-31+G(d) optimized geometries. Correlation effects were computed from the CCSD-MP2/6-31+G(d) energy difference [37].

The approach used in this work (B3LYP/aug-cc-pVTZ/B3LYP/6-31G(d)/SMD) performs well compared to the four approaches described above. For the total set of investigated tautomer pairs our approach has a RMSE of 2.2 [1.4,2.9] kcal/mol, making it the second best performing approach only outperformed by Kast et al. [15].

The difference in RMSE between the explanatory and blind dataset is noteworthy. Approaches that perform well on the blind data set perform worse on the explanatory set and vice versa. This is to a lesser extent also true for our chosen approach—B3LYP/aug-cc-pVTZ/B3LYP/6-31G(d)/SMD performs worse for the explanatory tautomer set (RMSE of 2.5 [1.5,3.6] kcal/mol) than on the blind tautomer set (RMSE of 1.3 [0.7,1.7]), but in comparison with the other four approaches, it is consistently the second best approach.

Interesting to note are the three tautomer pairs 15A_15B, 15A_15C and 15B_15C from the explanatory data set. The absolute error for these three pairs are 5.24, 1.9, and 2.8 kcal/mol, respectively. It appears that the used approach has difficulty to model 15B correctly, showing larger than average absolute errors whenever 15B is part of the tautomer reaction. Most likely the hydroxyl group in 15B is critically positioned and sensitive to partial solvent shielding by the phenyl ring, something that has been noted before [15].

The discrepancy between the different approaches (ours included) for the tautomer set shows that it seems highly difficult to propose a single method for different tautomer pairs that performs consistently with a RMSE below 2.0 kcal/mol. This issue is made substantially worse by the many different ways methods can be used/combined and errors can be propagated/compensated during tautomeric free energy difference 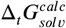 calculations. While we believe that a RMSE of 3.1 kcal/mol is a good value for the chosen approach, especially when compared to the results of the SAMPL2 challenge, it is by far not a satisfying result. The accuracy, compared to the cost of the approach, is not justifiable and there is still a dire need for more accurate and cheaper methods to obtain relative solvation free energies for tautomer pairs.

### Including multiple minimum conformations seems to have little effect on the accuracy of tautomer free energies

The three approaches described above consider multiple conformations to obtain the tautomeric free energy difference 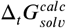. Obtaining the global minimum conformation is an important and well established task in quantum chemistry calculations. Figure 4 shows the difference between the highest and lowest free energy for an optimized minimum conformation for the individual tautomers of the dataset generated with B3LYP/aug-cc-pVTZ. While many of the tautomers have only a single minimum conformation (278 out of 936 tautomers), for molecules with multiple minimum conformations the difference in the free energy emphasizes the need and justifies the cost for a global minimum conformation search. Molecules with more than 10 kcal/mol difference between highest and lowest energy minimum are shown in Figure S.I.4.

While the scientific community agrees on the importance of the global minimum for property calculations, the importance of considering multiple conformations for quantum chemistry free energy calculations has not been well established (e.g. [38]). Using multiple minimum conformations can add substantial computational cost to the free energy calculations. Often, the global minimum conformation search can be performed with a lower level of theory than the single point energy calculation, making only a single high level electronic energy calculation necessary. Frequency calculations can also add considerably to the computational cost of the free energy calculation. If a single, minimum energy conformation is sufficient to obtain a good estimate for the free energy in gas phase a lot of computational time could be saved.

Figure 6 compares the tautomeric free energy difference obtained using the weighted average over multiple minimum conformations and the single global minimum conformation. The results indicate that using multiple minimum conformations does not significantly improve the final result compared to using a single, global minimum structure. Only 12 tautomer pairs show an absolute error larger than 1 kcal/mol in the tautomeric free energy difference 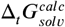 between the two approaches. Much more relevant than including multiple conformations is locating the global minimum conformation (as clearly shown by Figure 5).

**Figure 5.**
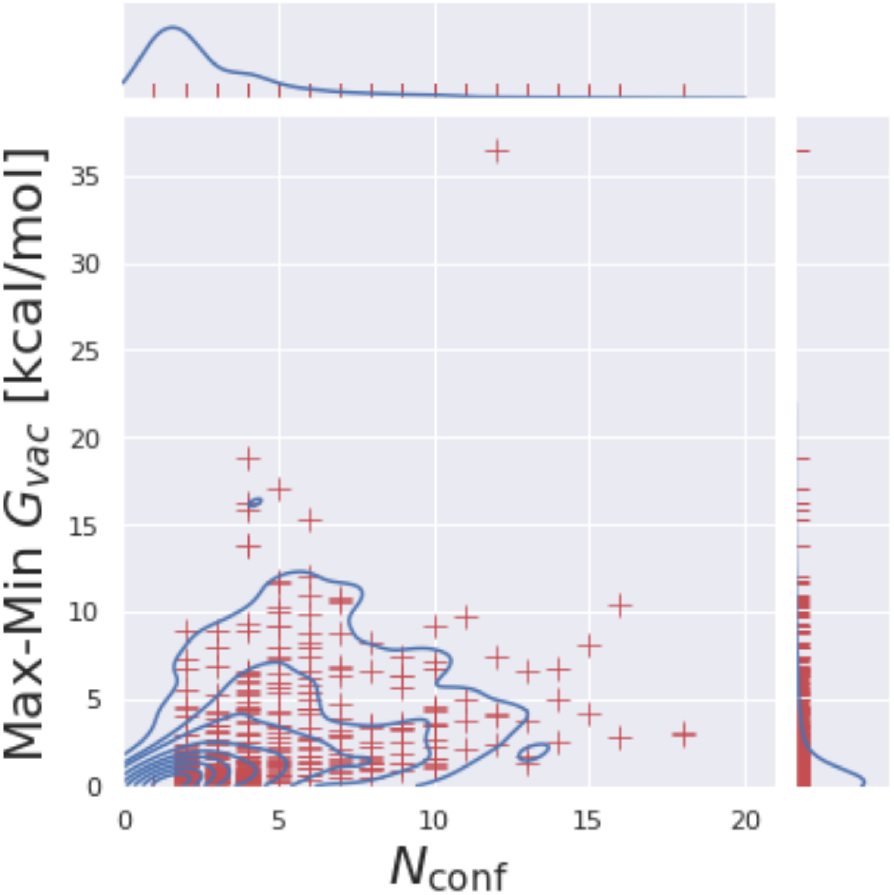
Computed free energies are highly sensitive to the selected minimum conformation. For each molecule, the number of minimum conformations *N*_conf_ is plotted against the difference between the corresponding highest and lowest obtained free energy value for the minimum conformations. 278 out of 936 molecules have only a single mini-mum; molecules with multiple minima show substantial free energy differences between the minimum conformations, highlighting the need for a global minimum search.

**Figure 6.**
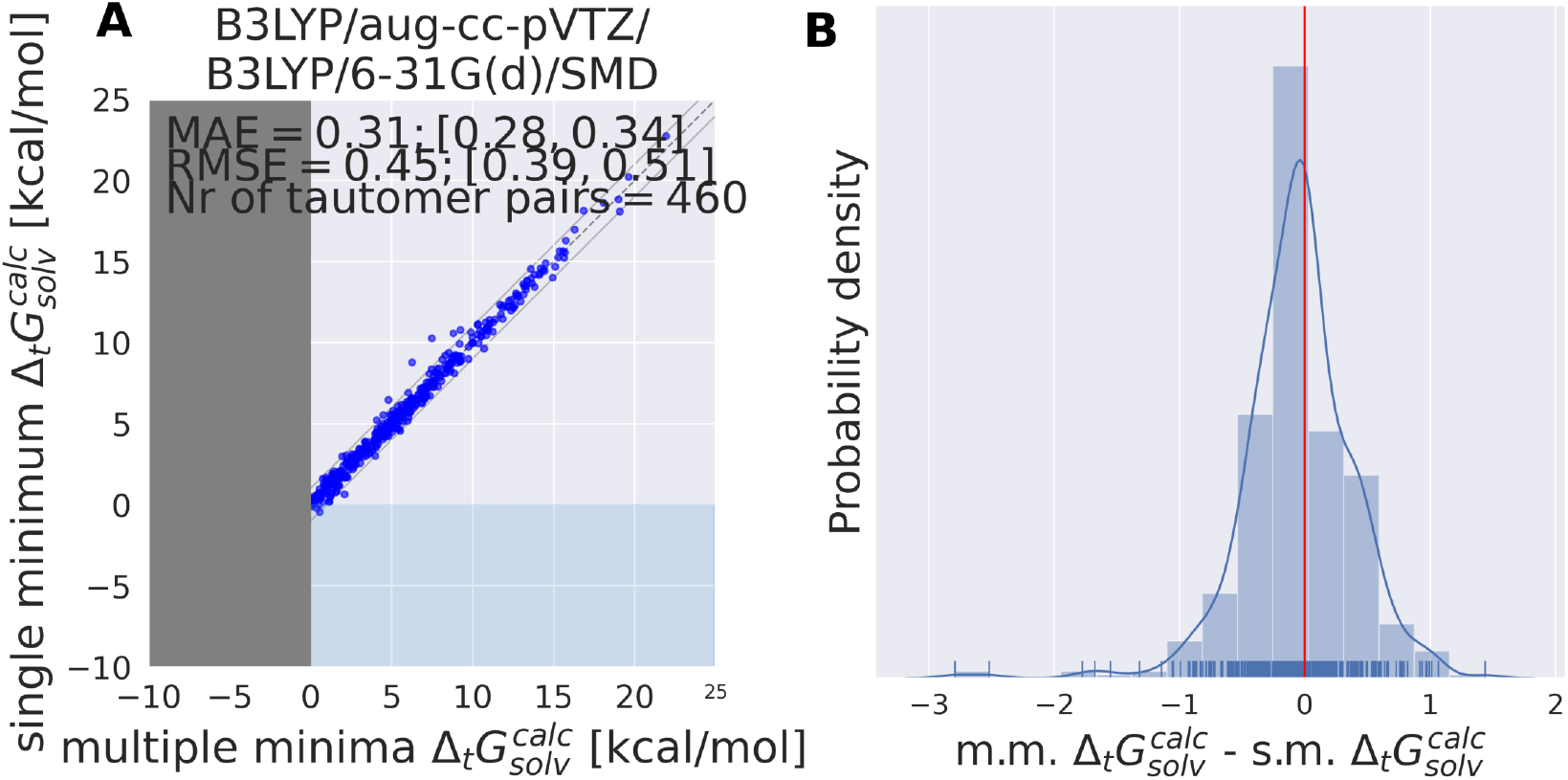
A single, global minimum conformation can be used to calculate tautomeric free energy differences in solution without loss of accuracy. These results are based on the calculations with B3LYP/aug-cc-pVTZ/B3LYP/6-31G(d)/SMD. **A** shows the tautomeric free energy difference 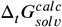 obtained with multiple minima (m.m.) plotted against the global, single minimum (s.m.) tautomeric free energy difference 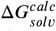. **B** shows the KDE and histogram of the difference between s.m. 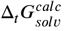 and m.m. 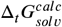. The comparison indicates that there is little benefit using multiple minimum structures over a single, global minimum considering the high costs of the former.

### Alchemical relative free energy calculations with quantum machine learning potentials can rigorously capture classical statistical mechanical effects

Alchemical relative free energy calculations were performed for 354 tautomer pairs using 11 alchemical intermediate λ states in vacuum.

In the following, we will compare the tautomeric free energy difference obtained using alchemical free energy calculations with the multiple minima, RRHO approximation using the same potential energy function (ANI-1ccx) to assess potential errors in the thermochemistry corrections. We will also show how a small number of experimentally obtained tautomer ratios in solution can be used to incorporate crucial solvent effects and recover tautomer free energies in solution by QML parameter optimization and importance weighting.

### RRHO ANI-1ccx calculations show significant deviations from the alchemical relative free energy calculations

In the limit of infinite sampling alchemical relative free energy calculations approach the exact free energy difference. Alchemical relative free energy calculations can be used to quantify the error introduced by a discrete partition function in the form of multiple minimum conformations and harmonic treatment of all bonded terms (including torsions and internal rotators)—if the same potential energy function is used for both calculations. Such a comparison is especially useful for quantum mechanics potentials which typically do not include hard-wired harmonic terms in their functional form (as is the case with most classical force fields).

Since the simulation time of the individual lambda states for the alchemical relative free energy calculations were relatively short (200 ps) we repeated the calculations multiple times (5) with randomly seeded starting conformations and velocities to detect systems for which the simulation time was clearly insufficient. In the following we will only use systems that had a standard deviation of less than 0.3 kcal/mol for 5 independent alchemical free energy calculations. Applying this filter resulted in the removal of 65 tautomer pairs for which the free energy calculation had not converged.

Results shown in Figure 7 indicate the average error that every relative free energy calculation based on the RRHO approximation introduces, regardless of the accuracy of the actual potential to model electronic energies. The mean absolute error of 0.9 kcal/mol should not be underestimated. Results shown in Table 1 would be significantly improved if an error of 0.9 kcal/mol could be compensated by running a protocol that samples the relevant conformational degrees of freedom to obtain an exact partition function.

**Figure 7.**
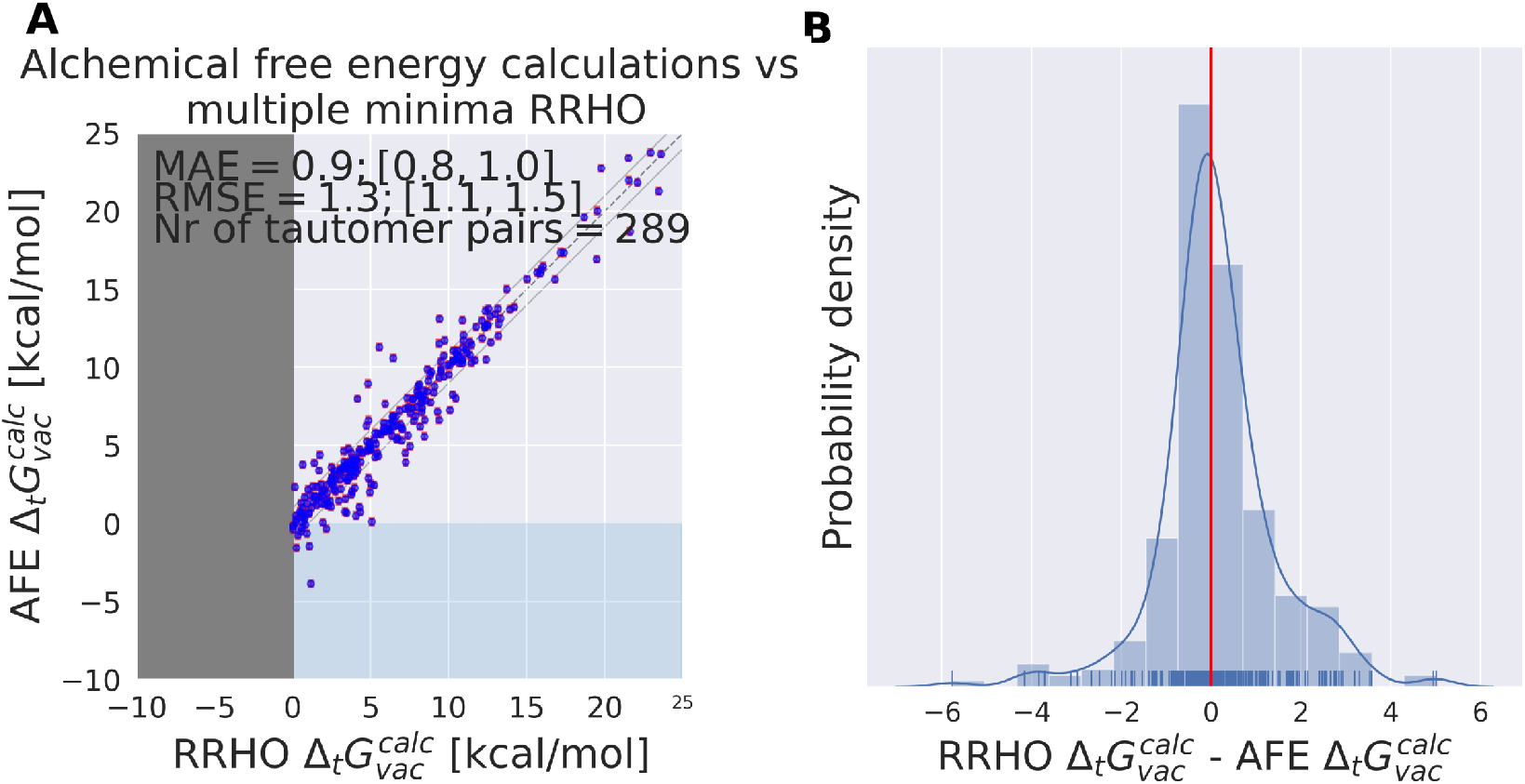
Independent of the level of theory, thermochemistry corrections introduce an average absolute error of 0.9 kcal/mol. ANI-1ccx is used for the calculation of tautomeric free energy differences in vacuum 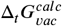 using alchemical relative free energy calculations and single point energy calculations on multiple minimum conformations and thermochemistry corrections based on the RRHO approximation (RRHO). **A** shows a scatter plot between the two approaches and **B** the KDE and histogram of the difference between the two approaches. Error bars are shown in red on the alchemical relative free energy estimates, these were obtained from the MBAR estimate of the relative free energy (see Methods section).

For 12 (out of 289) tautomer pairs the multiple minima RRHO approximation introduces errors of more than 3 kcal/mol (molecules are shown in Figure S.I.1). Most of these molecules have high conformational degrees of freedom and it seems unlikely that a naive enumeration of relevant conformations (e.g. with a conformer generator) will detect all of them — this might contribute to the observed error. That is certainly true for tp_113, 116, 565, 554, 403, 1674.

### QML potentials can be optimized to reproduce experimental tautomer ratios in solution

Since the alchemical free energy calculations were performed in vacuum, a comparison with the experimental tautomeric free energy difference 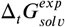 showed a high RMSE of 6.4 [5.9,6.8] kcal/mol, as expected from the well-known impact of solvation effects on tautomer ratios [39].

In the following we want to investigate if macroscopic properties — in this specific case tautomer ratios — can be used to retrain a neural net potential derived from quantum mechanics calculations. Further, we want to test how such an optimized parameter set would perform on the original dataset used to train the neural net potential and if it is possible to use the difference between the energies of the original and optimized parameter set for regularization.

Defining a loss function *L* as the error between the calculated 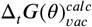 and experimental 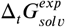 free energies it is possible to optimize the loss with respect to the neural net parameters *θ* defining the ANI potential. Using importance weighting, a new free energy estimate 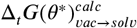 can be calculated with the optimized QML parameters (*θ*^*^) using the modified potential energy function without resampling the equilibrium distributions. To ensure that the quality of the neural net does not decrease beyond a reasonable threshold a regularization term that acts on the difference between the per snapshot energy obtained with the original and optimized parameter set Δ*E*(*θ, θ*^*^ was added. The regularization was included in the molecular loss function acting on the snapshots used for the free energy calculation for the 212 tautomer pairs in the training set — in the course of a single epoch Δ*E*(*θ, θ*^*)^ is evaluated on a total of 699,600 snapshots from 212 tautomer pairs, 11 lambda states and 300 snapshots drawn from each lambda state.

Initial results led to the introduction of scaling factors that allowed to slowly increase the weight of the contribution of the tautomeric free energy difference in the loss function (and/or the regularization term) during the training. The protocol is described in more details in the **Methods** section.

It was possible to overfit the parameters on the training set within 50 epochs if no regularization was used (training/validation performance shown in Figure S.I.5 I and M). Without regularization the RMSE for Δ*E*(*ρ, ρ*^*^) on the training/validation/test snapshots obtained from the MD simulations rises to 40-100 kcal/mol. While the training set performance improves in Figure S.I.5 I and M, the validation set performance reaches a plateau around 2.2–2.6 kcal/mol. The red dotted line in the Figure S.I.5 top panel indicates the performance of the reported results in Figure 8, indicating that even without regularization this performance can not be improved significantly. The results shown in Figure 8 and S.I.5 indicate that it is necessary to reduce the accuracy of the ANI1-ccx parameters on its original training set (either directly shown in Figure S.I.5 or indirectly through Δ*E*(*ρ, ρ*^*^) in Figure 8) to improve the performance for predicting tautomeric free energy differences. This is maybe best exemplified in the training/validation set performance shown in Figure S.I.5F and X. For both training runs the scaling factor of the regularization term was increased throughout the epochs, shifting the focus from optimizing the free energy to keeping Δ*E*(*ρ, ρ*^*^) as small as possible. In Figure S.I.5 X the Δ*E*(*ρ, ρ*^*^) term in the loss was kept constant throughout the first 100 epochs and then raised from 3 to 25 exponentially during the next 50 epochs. Looking at the training/validation set performance it becomes evident that starting with epoch 100 the initial gained better performance starts to vanish while the MAE for Δ*E*(*ρ, ρ*^*^) decreases. After 400 epochs the MAE for Δ*E*(*ρ, ρ*^*^) is near zero and there is no improvement in the free energy estimate for the training/validation set. This indicates that we are approaching the same minimum that was occupied before training started. Whenever both weights and biases are trained it often takes a few hundred epochs before Δ*E*(*ρ, ρ*^*^) approaches reasonable values (∼ 2 kcal/mol), shown e.g. in Figure S.I.5 L, J, G, and A. This also happened when the learning rate for the biases was reduced to 1e-6 as shown in Figure S.I.5T. In some instances it was not possible to converge and successfully minimize both terms of the loss function in 400 epochs. One such case is shown in Figure S.I.5E.

**Figure 8.**
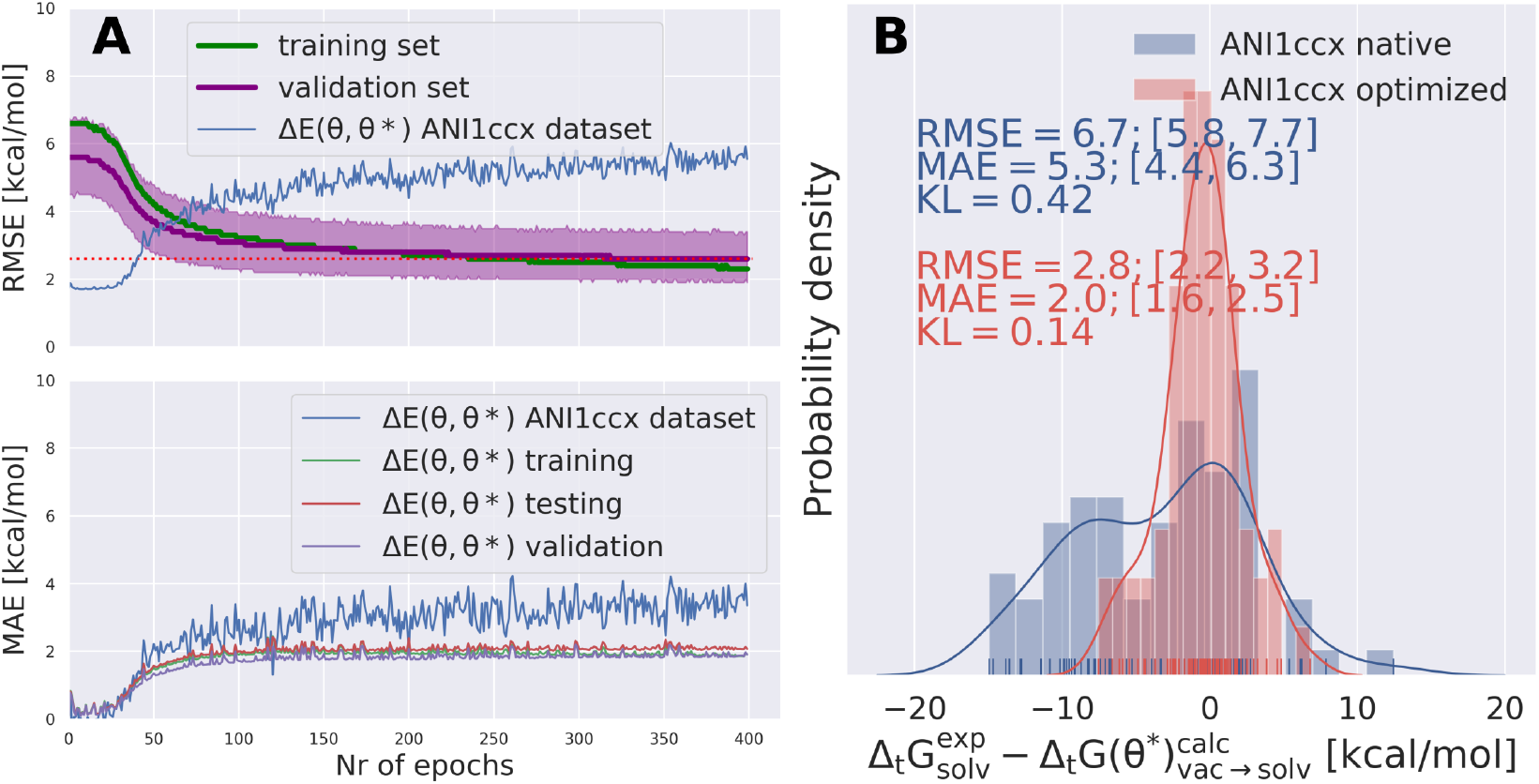
Optimizing QML parameters on a set of experimentally obtained tautomer free energies in solution 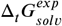 enables ANI-1ccx to include crucial solvation effects and improved estimates for tautomeric free energy differences can be obtained by importance weighting from vacuum simulations using the optimized QML parameters. **A** top panel shows the training (green) and validation (purple) set performance (RMSE) as a function of epochs. Validation set performance was plotted with a bootstrapped 95% confidence interval. The performance of the optimized parameter set is also shown on the original ANI-1ccx dataset. The best performing parameter set (evaluated on the validation set – indicated as the intersection between the two red dotted lines) was selected to compute the RMSE on the test set. The bottom panel shows the MAE for the per snapshots energy difference between each of the 400 parameter sets and the original parameter set on all the snapshots used for the free energy calculations (1,2 million snapshots) split in training/validation and test set as well as the original ANI1-ccx dataset. Figure **B** shows the distribution of 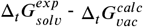 for the tautomer test set (71 tautomer pairs) with the native ANI-1ccx (*θ*) and the optimization parameter set (*θ*)*. The RMSE of tautomeric free energy difference was improved from 6.7 kcal/mol to 2.8 kcal/mol (MAE improved from 5.3 to 2.0 kcal/mol) by optimizing the parameter set. The difference in Kullback-Leibler divergence (KL) indicates that the tautomeric free energy differences obtained with the optimized parameter set (0.14) can reproduce the distribution of the experimental tautomer ratios much better than the the free energy differences obtained with the original parameter set (0.42).

In Figure 8 A the training/validation set performance of a training run with 400 epochs is shown. nly weights were optimized with a learning rate of 1e-4, regularization was used (the scaling factors are plotted in Figure S.I.6). Figure 8 B shows the obtained free energy estimates 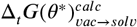 with the best performing parameters set (calculated on the validation set) on an independent test set (70 tautomer pairs). The RMSE on the test set after optimization was decreased from 6.7 [5.7,7.7] kcal/mol with the original parameter (*θ*) set to 2.8 [2.2,3.2] kcal/mol with the optimized parameter set (*θ*^*^), comparable to the performance of B3LYP/aug-cc-pVTZ/B3LYP/6-31G(d)/SMD. Figure 8 B shows the distribution of 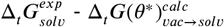. The Kullback-Leibler divergence (KL) value for 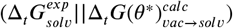 indicates that overall the optimized parameter set is able to reproduce the distribution of the experimental free energies better.

Figure 8 A shows the performance of the parameter set on the ANI-1ccx training dataset^1^. The RMSE of the parameter set for the ∼ 500k data points of the dataset increases from initial 1.7 kcal/mol to 5.5 kcal/mol. This increase in RMSE can be partially explained by the target of the retraining: tautomeric free energies *in solution*. Including solvation effects necessarily decreases the performance on a purely gas phase

Coming back to the question posed in the beginning of this section: yes, it appears to be possible to use experimental values of a thermodynamic observable to retrain a already optimised QML potential to better perform on the task of predicting this macroscopic property. This is astonishing, considering the very small set of experimental measurements used to retrain the potential (≈ 250 data points) that provides an unambiguous and substiantial improvement in performance on the held-out test set. The regularization term restraining the QML parameter set so that it is still able to reproduce single point energies with comparable quality than the original parameter set (with an increase in RMSE of ∼ 3 kcal/mol) offers an interesting trade off between high quality QM single point energies and highly improved tautomeric free energy differences. It seems that the regularization is not necessary to improve the tautomeric free energy difference estimate, but introducing such a regularization term in the loss function does also not significantly hinder the parameter set optimization (if the scaling factors for the two terms in the loss function are chosen reasonably). These results highlight the incredible potential of QML energy functions that are easily differentiable with respect to input parameters in addition to coordinates, since this enables model tuning based on experimental and quantum chemical data to be both facile and incredibly powerful.

## Discussion & Conclusion

In this work we use a state of the art density functional theory protocol and continuum solvation model to calculate tautomer ratios for 460 tautomer pairs using three different approaches to model the solvent contributions. The best performing method uses B3LYP/aug-cc-pVTZ and the rigid rotor harmonic oscillator approximation for the gas phase free energies (calculated on B3LYP/aug-cc-pVTZ optimized geometries). The transfer free energy was calculated using B3LYP/6-31G(d)/SMD on geometries optimized in their respective phase (with B3LYP/aug-cc-pVTZ). This approach performs with a RMSE of 3.1 kcal/mol.

One possible source of error — independent of the method used to calculate the electronic energy and model the continuum electrostatics — are the thermochemical corrections used to obtain the standard state free energy. Typically, an analytic expression is used to approximate the partition function which is based on the rigid rotor harmonic oscillator approximation. To obtain the correct and unbiased partition function and compute rigorous free energy estimates we implemented an alchemical relative free energy workflow using the ANI family of quantum machine learning (QML) potentials [24]. The method was implemented as a python package and is available here: https://github.com/choderalab/neutromeratio.

Using the same potential for the calculations based on the RRHO and performing alchemical relative free energy calculations we are able to show that the RRHO approximation introduces a mean absolute error of ∼ 1 kcal/mol in the calculatino of the investigated tautomeric free energy differences. These errors can be attributed to anharmonicity in bonded terms, difficulties to enumerate relevant minimum conformations and in combining shallow local energy wells as well as the inconsistent treatment of internal and external symmetry numbers.

The calculated alchemical free energies obtained using the methods implemented in the “‘Neutromeratio”’ package can be used to optimized the parameters of the QML potential. Using a small set of experimentally obtained tautomeric free energy differences we were able to optimize the parameter set on a training set and significantly improve the accuracy of the calculated free energies on an independent test set. The tautomeric free energies obtained with the optimized parameter set had a RMSE of 2.8 kcal/mol.

What should be noted here: the experimental values are relative solvation free energies, while we calculate relative gas phase free energies. The optimization routine on the experimental relative solvation free energies adds crucial solvent effects to the accurate description of the vacuum potential energy. The ANI family of QML and comparable QML potentials have opened the possibility to investigate tautomer ratios using relative free energy calculations without prohibitive expensive MM/QM schemes.

Obtaining accurate free energy differences between tautomer pairs in solvent remains an elusive task. The sublet changes and typically small difference in internal energies between tautomer pairs require an accurate description of electronic structures. Furthermore, solvent effects have a substantial effect on tautomer ratios; consequently, a proper descriptor of solvation is essential. The change in double bond pattern typically also induce a change in the conformational degrees of freedom and, related, in the conformation and rotational entropy. But — despite all of these challenges — we remain optimistic that further developments in fast and accurate neural net potentials will enable improved and more robust protocols to use relative free energy calculations to address these issues.

## Code and data availability

- Python package used in this code (release v0.2): https://github.com/choderalab/neutromeratio
- Data and notebooks to reproduce the plots/figures (release v0.2): https://zenodo.org/record/4562426

## Author Contributions

Conceptualization: JDC, JF, and MW; Methodology: JDC, JF, and MW; Software: JF, and MW; Investigation: JF, and MW; Writing–Original Draft: JDC, JF, and MW; Writing–Review&Editing: JDC, JF, and MW; Funding Acquisition: JDC and MW; Resources: JDC and MW; Supervision: JDC and MW.

## Acknowledgments

MW, JF and JDC are grateful for discussions from the Tautomer Consortium, specifically Paul Czodrowski, Brian Radak, Woody Sherman, David Mobley, Christopher Bayly and Stefan Kast as well as input from Adrian Roitberg and Olexandr Isayev. This work was only possible through the time and effort Thomas Sander and Oya Wahl invested in curating the open tautomer database (Tautobase). Further, MW is grateful for the support of Thierry Langer, who donated significant computational resources to perform much of the calculations/simulations and Oliver Wieder, for helpful discussions.

## Detailed methods

### Experimental data

The full dataset considered for this study was obtained from the DataWarrior File deposited in https://github.com/WahlOya/Tautobase (commit of Jul 23, 2019), described in detail in [10]. The dataset was sourced from the tautomer codex authored by P.W. Kenny and P.J. Taylor [35].

From the dataset a subset of tautomer pairs were considered that

1. were measured/calculated/estimated in aqueous solution
2. had a numeric logK value between +/-10
3. had no charged species
4. did not contain iodine
5. only a single hydrogen and double bond change its position

476 of the 1680 deposited tautomer pairs had these properties. We added two tautomer pairs from the SAMPL2 challenge (Tautomer pair 2A_2B and 4A_4B) [13]. 478 unique tautomer pairs were considered for further analysis. The term ‘unique’ tautomer pair was defined in the following as containing a unique combination of two molecules. In the following we will use an identifier containing the row number entry from the DataWarrier file to identify tautomer pairs in the dataset, e.g. tp_200 describing the tautomer pair (tp) at row number 200 in the original DataWarrior file.

The logK value was converted to free energies with 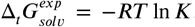. The free energy difference of the tautomer pairs obtained from the Tautobase are subsequently referenced as 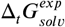 in contrast to the calculated values which are called Δ_*t*_*G*^*calc*^. While we call all values deposited in the Tautobase 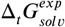 we want to point out that some of these values are estimated or calculated.

A closer inspection of some of the outliers from preliminary calculations identified molecules with incorrect structures in the database (Row entry: 1260,1261,1262,1263,1264,514,515,516,517,1587) — these 10 tautomer pairs were subsequently removed from all QM calculations (simulations using ANI1 included the corrected tp_1260, tp_1261 and tp_1587). These structures are corrected in the current version of the Tautobase. Additionally, 8 tautomer pairs containing bromide (Row entry: 989, 581, 582, 617, 618, 83, 952, 988) were removed from calculations performed with the basis set 6-31G(d) due to the lack of adequate parameters.

ANI1 is parameterized for four elements Carbon, Nitrogen, Hydrogen and Oxygen, making a further removal of tautomer pairs containing elements beside the aforementioned [24]. This resulted in 369 tautomer pairs.

The tautomer pair set used for relative alchemical free energy calculations needed one additional filter. Molecules with a stereobond that changes its position between tautomers had to be removed (this affected 15 tautomer pairs: tp_1637, tp_510, tp_513, tp_515, tp_517, tp_518, tp_787, tp_788, tp_789, tp_810, tp_8Δ, tp_812, tp_865, tp_866, tp_867). Such stereobonds introduce additional complexity since it would be necessary to introduce restraints to define the stereochemistry during the lambda protocol.

The subset of the Tautobase used for the QM calculations can be obtained here as list of SMILES (468 tautomer pairs): https://github.com/choderalab/neutromeratio/blob/master/data/b3lyp_tautobase_subset.txt. The subset of the Tautobase used for the QML calculations can be found here as list of SMILES (354 tautomer pairs): https://github.com/choderalab/neutromeratio/blob/master/data/ani_tautobase_subset.txt. The distribution of the experimental tautomer free energies for both datasets is shown in Figure 9.

### Generating molecular conformations

The input tautomer pairs were specified as SMILES strings. 3D conformations were generated with the chemoinformatics toolkit RDKit (version 2019.09.2) which uses the distance geometry approach to generate conformations while enforcing chirality/stereochemistry [40, 41] For each molecule 20 conformations were initially generated. The number of conformations was reduced to 10 if the average root mean square deviation (RMSD) of atoms of the 20 conformations was below 0.5 Angstrom, and further reduced to 5 conformations if below 0.2 Angstrom.

**Appendix 0 Figure 9.**
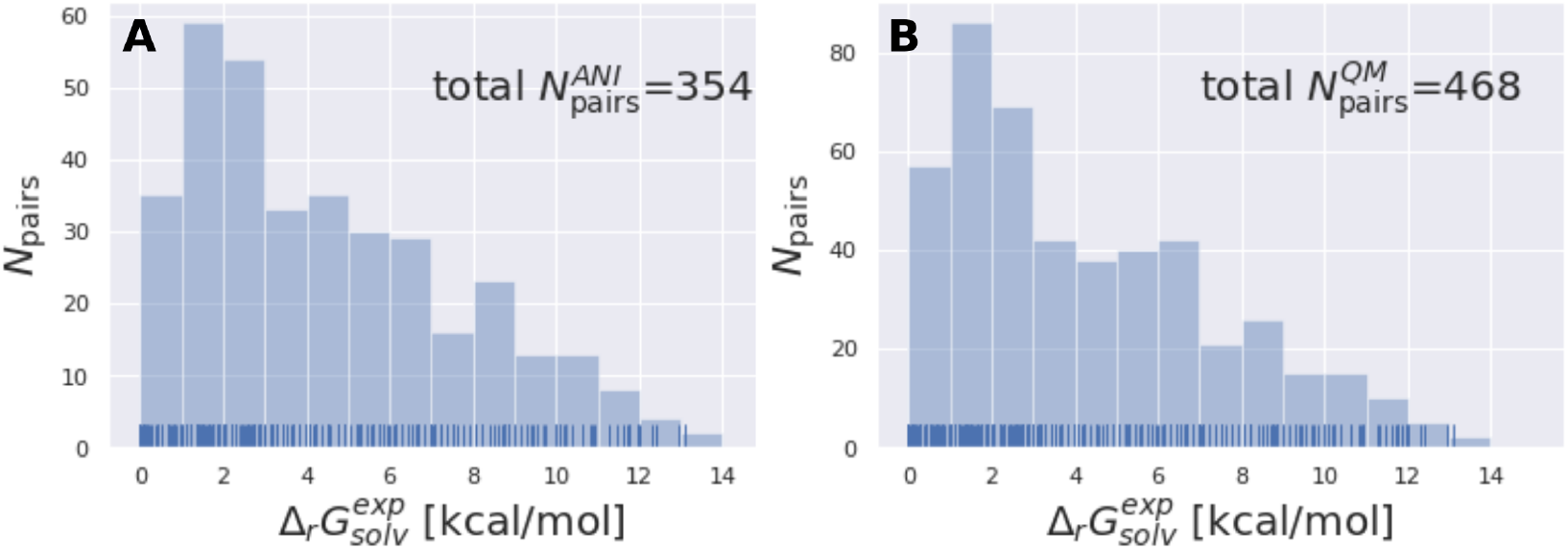
The tautomer dataset shows a wide variety of solvation free energies. 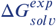. **A** shows the ANI-Tautobase subset that was used for the QML calculations and **B** shows the QM-Tautobase subset used for the QM calculations. The full dataset considered for this study was obtained from the DataWorrier File deposited at https://github.com/WahlOya/Tautobase(commit of Jul 23, 2019), described in detail in [10]. The selection criteria for both datasets are described in detail in Detailed methods section.

### RMSD calculations and filtering of conformations

For each molecule, pairwise RMSD between conformations were calculated using RDKit. Starting with a random conformation, if the RMSD to any other conformation of the molecule was below 0.1 the conformation is discarded, otherwise added to the list of unique conformations. The RMSD was calculated between heavy atoms and the hydrogen of selected chemical moieties including primary alcohols, imines, primary/secondary amines, cyanamides and thiols.

### Combining energies of conformations

Energies of different minimum conformations of the same molecule were weighted and combined using the following scheme

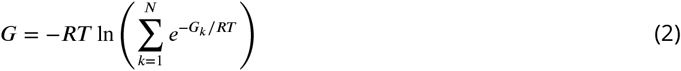

The obtained free energies will be called weighted free energies in the following.

### Quantum mechanical calculations

The following quantum mechanical (QM) calculations were performed using the quantum chemical software orca 4.0.1.2 [42]. The universal solvation model based on solute electron density (SMD) was used as continuum solvation model [33].

#### Geometric optimization, single point energy and frequency calculation

Geometric optimization was performed with the standard options of orca, redundant internal coordinates and the BFGS optimizer [28–30]. Frequency calculations were performed with the numerical Hessian computed using the central differences approach. If a conformation had negative frequencies (imaginary modes) after the geometry optimization it was excluded from further analysis. Single point calculations were performed on the optimized geometries using B3LYP and the basis set aug-cc-pVTZ or 6-31G(d) [31, 32]. A damping dispersion correction was applied (orca keyword D3BJ) [43].

#### Continuum solvation model

The continuum solvation model SMD was used to model the molecules in aqueous environment [33]. Since there is a volume change in the standard state from the gas phase (1 atm) to the solvent phase the gas phase standard state is indicated by ‘*’ and the solvation standard state by ‘o’. The standard-state transfer free energy is then defined as

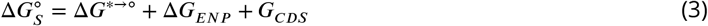

with Δ*G*^*→o^ as standard-state adjustments (specifically, the correction of changing the volume from the gas phase to the solute phase with a constant value of 1.89 kcal/mol), Δ*G*_*ENP*_ describes the electronic (E), nuclear (N) and polarization (P) components of the free energy and Δ_*CDs*_ free energy changes associated with solvent cavitation (C), changes in dispersion (D) and changes in local solvent structure (S) [33].

#### Free energy and free energy in solution calculations

Thermal corrections were computed at standard state (298.15 K and 1 atm pressure) using the ideal gas molecular partition function and the rigid-rotor harmonic oscillator (RRHO) approximation. Low-lying vibrational frequencies (below 15 cm^−1^) were treated by a free-rotor approximation [44] — this method is also sometimes called rigid-rotor quasi harmonic oscillator. The external (rotational) symmetry number was obtained from the point group of the tautomer using the point group module of Jmol and visual inspection and used to correct the rotational entropy calculated by orca [45].

The Gibbs free energy in gas phase 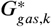 for a given coordinate set (*k*) was obtained by adding thermal corrections 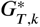 for conformer *k* at temperature *T* and zero point energy (ZPE) contributions ϵ_ZPE,*K*_) to the electronic energy *E*_*k*_ [23]. The degeneracy *D* describes the entropy contribution of internal rotors and is not added for the calculations shown in the main text (results including degeneracy *D* are shown in the Supplementary Material).

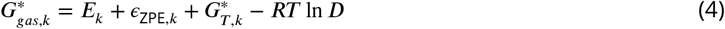

The degeneracy *D* was estimated by calculating the graph automorphism of the molecule. The implementation of the VF2 algorithm for graph isomorphism of networkx was used [46]. Nodes were defined to match if element and hybridization matched, edges were identical if bond order matched.

The tautomeric free energy difference in solution 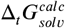 can be calculated from the standard-state Gibbs free energy in aqueous phase 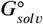 of the product and educt of the corresponding tautomer reaction, which itself is calculated as the sum of the gas-phase standard-state free energy 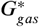 and the standard-state transfer free energy 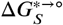. expressed as

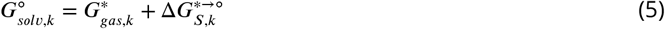

for a given conformation *k* and shown as a thermodynamic cycle in Figure 1.

An alternative way to calculate the free energy in aqueous phase 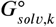 is

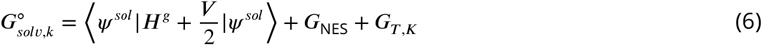

Where *ψ* ^*sol*^ is the polarized wave function in solution, *H*^*g*^ the gas phase Hamiltonian, and V the potential energy operator associated with the reaction field. The bracket term describes the electronic energy, while *G*_NES_ is associated with non-electrostatic contributions (dispersion-repulsion and solvent structural terms) to the solvation energy and *G*_*T,K*_ are the thermal correction calculated directly in the continuum solvation model [47].

### ASE thermochemistry corrections

ANI-1ccx was used to calculate the electronic energy. The conformations were minimized using the BFGS optimizer as implemented in scipy [48]. Frequency and thermochemistry calculations were performed using the optimized geometry. Thermal corrections were calculated for 298 K and 1 atm using the IG-RRHO approximation as implemented in the atomic simulation environment (ase) [49]. The tautomeric free energy difference in gas phase Δ_*t*_*G*_*vac*_ was then calculated as the difference between 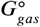 of the tautomer pair.

### Relative alchemical free energy calculations

Relative alchemical free energies were calculated using a single, hybrid topology approach. The topology of each tautomer pair only differed in the position of a single hydrogen (atom types and bonds were not specified). The hybrid topology (the superset of the two topologies) differed therefore by one hydrogen from each of the physical endstates.

By default, the coordinates of the hybrid topology were generated by using the coordinates of ‘Tautomer1’ (as defined in the Tautobase database). If a tautomer isomerismn created or removed a cis/trans stereobond, the initial coordinates were taken from the topology with the stereobond present (therefore sometimes changing the direction of the tautomer reaction).

The coordinates of the added, non-interacting hydrogen were obtained by randomly sampling 100 positions on the surface of a sphere (with radius of 1.02 Angstrom) defined around its new bonded heavy atom and subsequently using the lowest energy position. The physical endstates (representing the two tautomer states, each with an additional non-interacting (dummy) hydrogen) were connected via Δ intermediate (lambda) states.

Energy and forces were calculated using ANI1-ccx as implemented in torchani https://github.com/aiqm/torchani). The energy and force was linearly scaled along the alchemical path as a function of lambda with

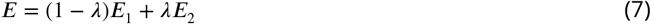

Before each simulation, initial coordinates were minimized using a BFGS optimizer as implemented in scipy [48]. Coordinates were sampled using Langevin dynamics at 300 K with a collision rate of 10 ps^−1^ and a 0.5 fs time step using the BAOAB integrator [50]. Initial velocities were obtained from a Maxwell-Boltzmann distribution at the simulation temperature.

Samples were obtained from 200 ps simulations for each lambda state. For each tautomer pair calculations were repeated 5 times with randomly seeded initial velocities (and coordinates).

All bonds involving hydrogen were restrained throughout the simulation at each lambda states using a flat bottom potential well with harmonic walls. The restraint was defined as

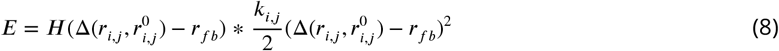

With *H* as the Heaviside step function, 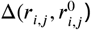 as the difference between the reference bond length 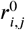 and the current bond length *r*_*i,j*_ and *r*_*fb*_ as half of the well radius. For all heavy atom pairs 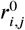 was set to 1.3 Angstrom and *r*_*fb*_ to 0.3 Angstrom. For C-H/O-H/N-H bond pairs 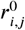 was set to 1.02 Angstrom (the average of the three different equilibrium bond length of C-H/O-H/N-H bond pairs) and *r*_*fb*_ to 0.4 Angstrom.

Relative alchemical free energies 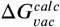 were calculated using MBAR as implemented in the pymbar package [51]. 300 uncorrelated snapshots were considered for the MBAR analysis from each lambda state.

### Neural net parameter optimization based on experimental relative solvation free energies

The tautomer data set was randomly split (20:20:60) into a test set (71), validation set (71) and training set (212 tautomer pairs). Neural net parameters were optimized using a routine modified from the TorchANI tutorial^2^. To limit the capacity to overfit, only the weights and biases of the *final* layer of each of the 8 pretrained ANI-1ccx models were optimized for each of the atom nets (one net per element), resulting in roughly 8×4×97 tunable parameters (8 neural nets, each with 4 atom nets containing 97 weights and bias) for ANI-1ccx. As in the TorchANI tutorial, the weight matrices were updated using the Adam optimizer with decoupled weight decay (AdamW), and the bias vectors were updated using Stochastic Gradient Descent (SGD). The training data was randomly partitioned in each epoch in mini-batches of 10 tautomer pairs and gradient updates were performed for each mini-batch. Training was performed for 400 epochs. The best model was chosen based on the RMSE on the validation set and model performance reported on the test set.

The model was trained by minimizing the mean squared error (MSE) loss between calculated and experimental relative free energies. If regularization was used, the mean absolute error (MAE) between the energy for the parameter set *θ*^*^ and the original parameter set *θ* was calculated on the individual snapshots used for the free energy calculation (Δx300 snapshots). The regularization term was normalized using the number of atoms of the tautomer system. The per molecule pair (*m*) loss function is defined as

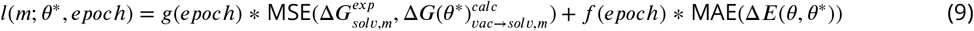

with 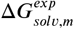 as the experimental and 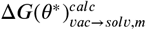 as the calculated tautomeric free energy differences for molecule m using parameters *θ*^*^. The two scaling factors *f* (epoch) and *g*(epoch) were used to control the contribution of the two terms as a function of training epoch. For the final results the values for g (labeled ‘scaling dG’) and f (labeled ‘scaling dE’) is shown in Figure S.I.6.

The overall loss is then

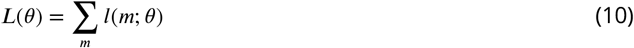

The perturbed relative free energy 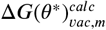 required for computing (*m*; *θ*^*^) is calculated by importance sampling, or “reweighting” the original MBAR estimate from the original pre-trained parameters *θ* to the current parameters *θ*^*^. To compute this estimate efficiently at arbitrary *θ*^*^, we first collect configuration samples at a reference value *θ* (corresponding to the original parameters of the pretrained ANI-1ccx model) for each intermediate value *λc*. For each configuration sample *x*, we compute the reduced potential *u*(*x, λc*; *θ*), to form the *N* × *M* matrix of inputs to MBAR. (Where *N* is the total number of snapshots, *M* is number of *λ* windows.) MBAR equations are solved to yield a vector of reduced free energies 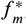, where *m* indexes the intermediate *λ* values. The relative free energy prediction implied by the model parameters *θ*^*^ is then 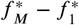.

To compute relative free energies at a new value of the parameters *θ*^*^, we need to compute *u*(*x, λc* = 1; *θ*^*^) and *u*(*x, λc* = 0; *θ*^*^) for all configurations *x*. The optimization routine evaluates the gradient of the loss function *L* w.r.t. *θ*^*^ by automatic differentiation and updates the parameters.

300 uncorrelated snapshots per *λc* state were used for the MBAR estimate and importance sampling for the vacuum simulations.

**Appendix 0 Figure S.I.1.**
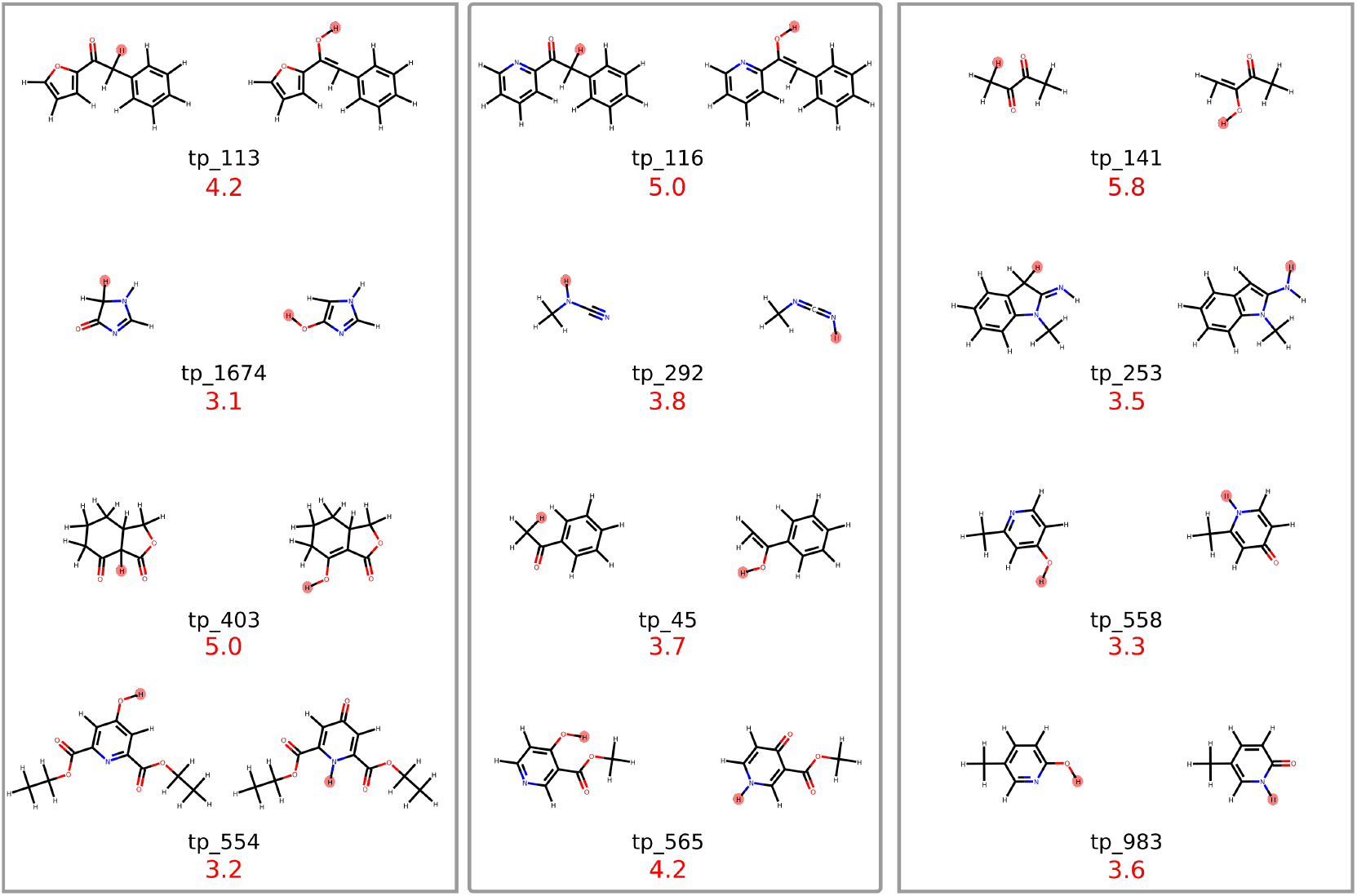
Molecules for which the RRHO approximation introduces an error of more than 3 kcal/mol are shown. Absolute error is shown in red (value in kcal/mol) The hydrogen that changes position is high-lighted in red.

## Supplementary Information

**Appendix 0 Table S.I.1.**
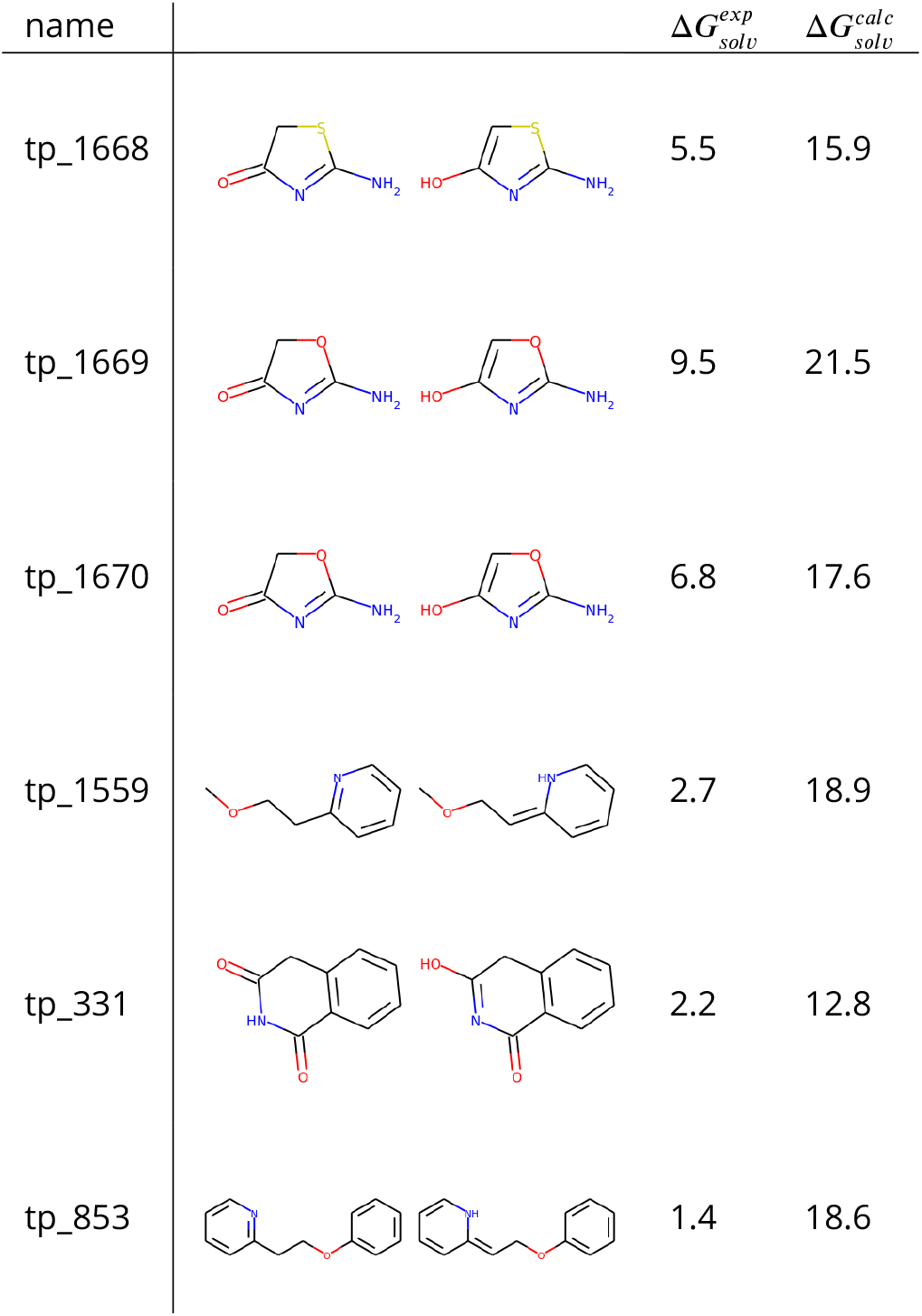
5 out of the 6 tautomer pairs with the highest absolute error have common scaffolds. Tautomer pairs with absolute errors above 10 kcal/mol are shown.

| name | | $\Delta G_{solv}^{exp}$ | $\Delta G_{solv}^{calc}$ |
| --- | --- | --- | --- |
| tp_1668 |  | 5.5 | 15.9 |
| tp_1669 |  | 9.5 | 21.5 |
| tp_1670 |  | 6.8 | 17.6 |
| tp_1559 |  | 2.7 | 18.9 |
| tp_331 |  | 2.2 | 12.8 |
| tp_853 |  | 1.4 | 18.6 |

**Appendix 0 Figure S.I.2.**
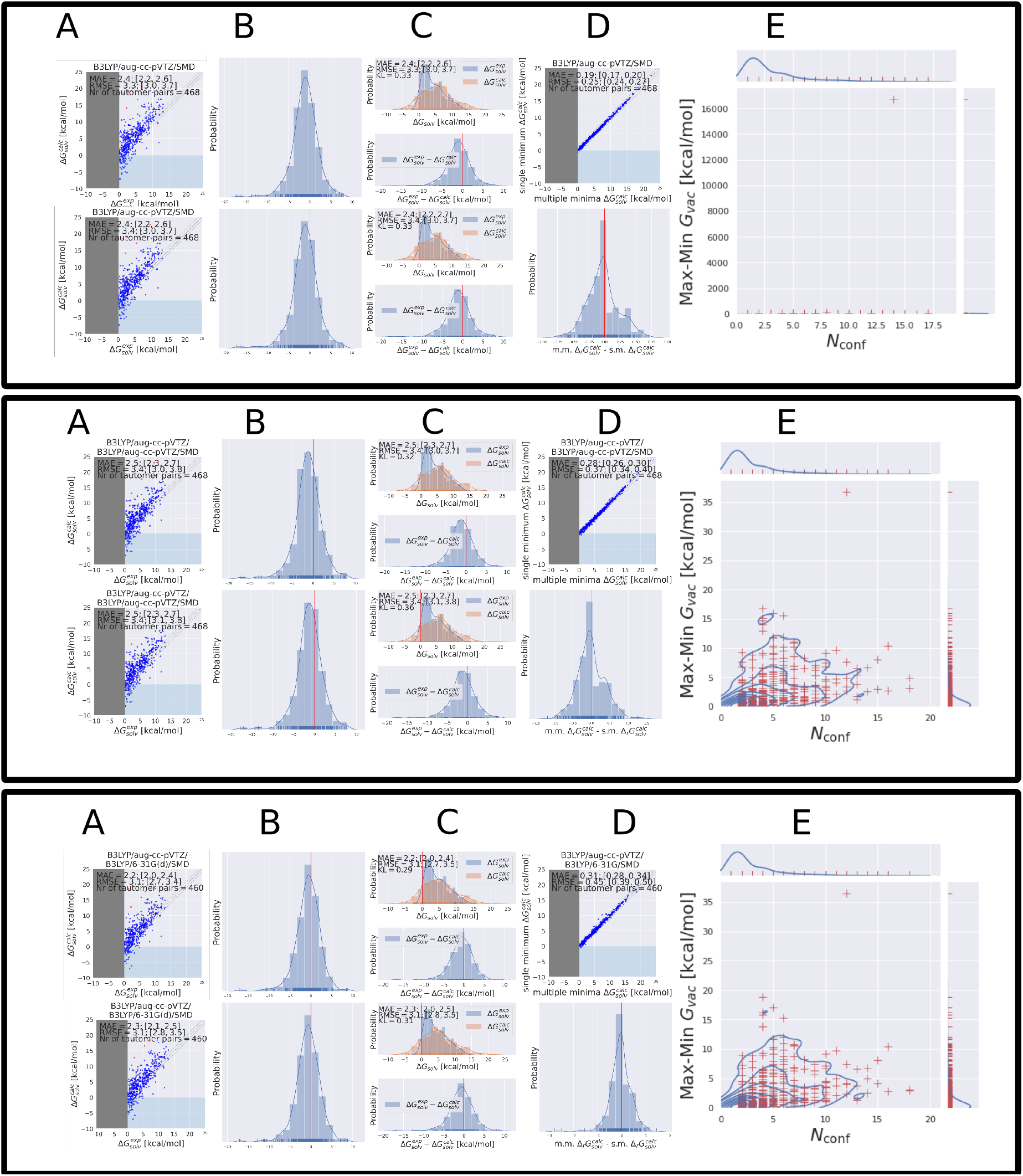
Each of the three blocks show the QM calculations done with different level of theory indicated by the title in the plots in column **A**. In each block the top penal shows the results obtained with multiple minimum conformations, the bottom penal shows the results obtain with a single conformation. Column **D** shows the difference between the multiple minimum and single minimum approach and **E** show the the number of minimum conformations against the difference between the lowest and highest energy for each tautomer. These results were obtained without the additional structure symmetry correction.

**Appendix 0 Figure S.I.3.**
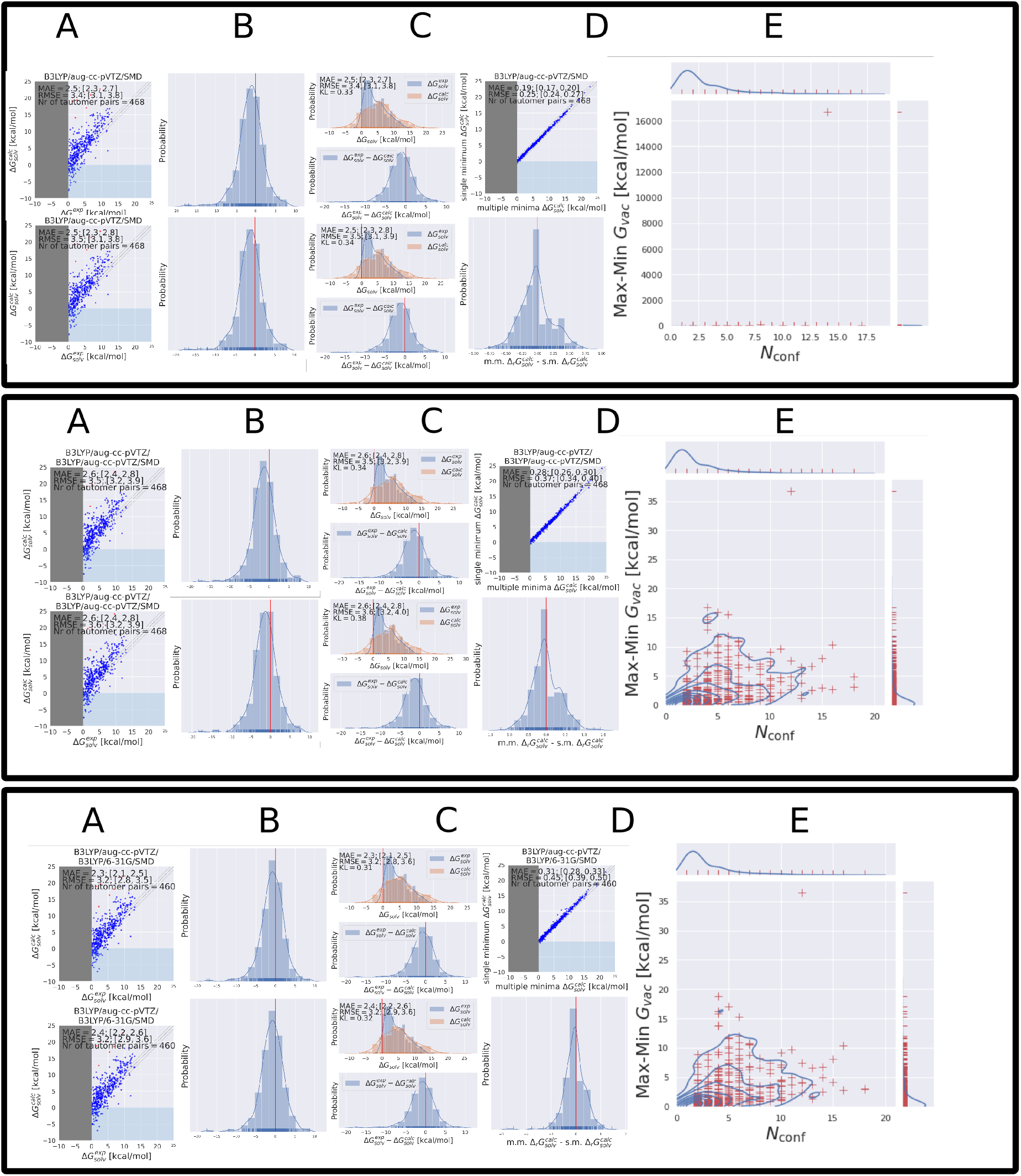
Each of the three blocks show the QM calculations done with different level of theory indicated by the title in the plots in column **A**. In each block the top penal shows the results obtained with multiple minimum conformations, the bottom penal shows the results obtain with a single conformation. Column **D** shows the difference between the multiple minimum and single minimum approach and **E** show the the number of minimum conformations against the difference between the lowest and highest energy for each tautomer. These results were obtained **with** structure symmetry correction.

**Appendix 0 Figure S.I.4.**
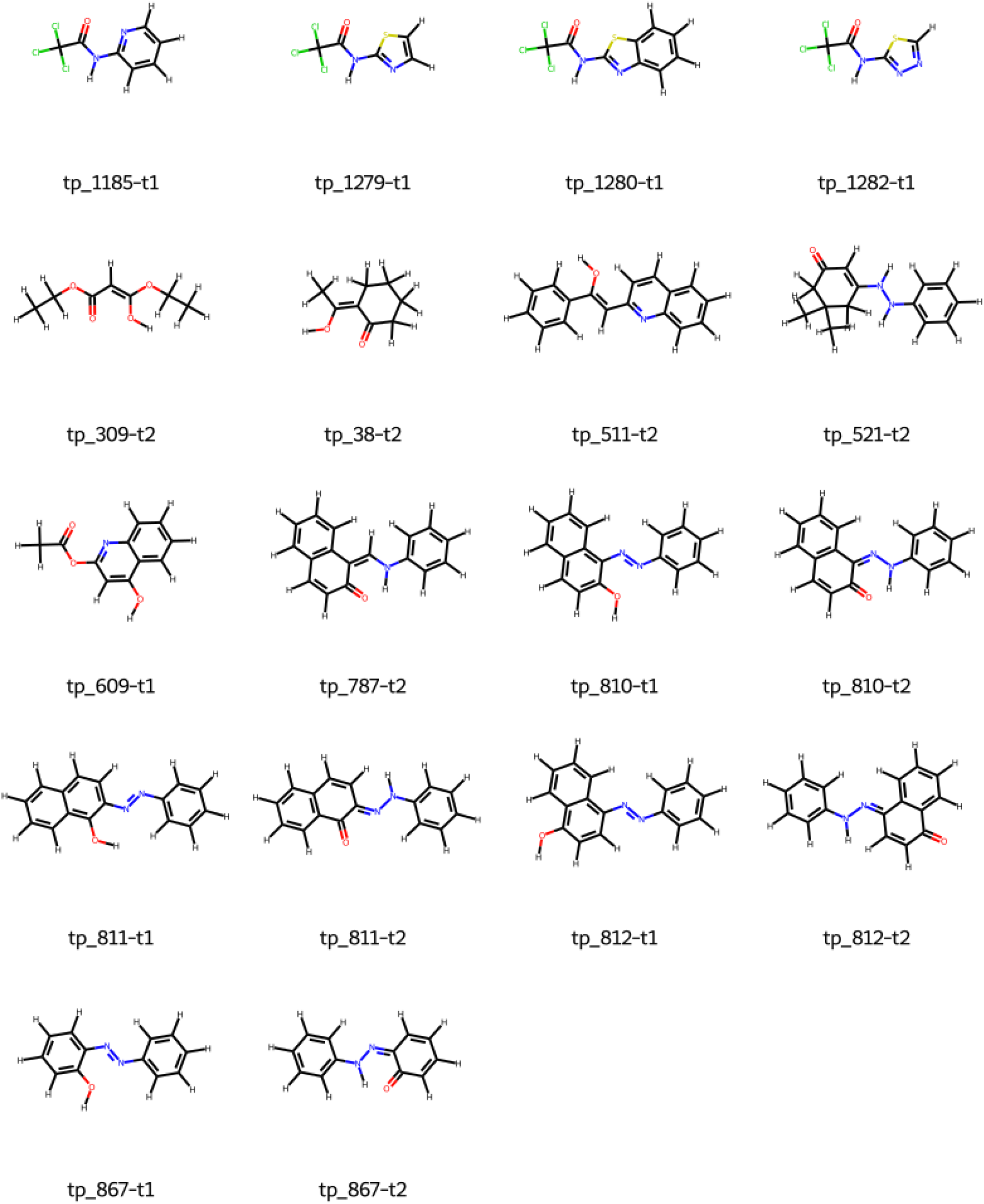
Molecules for which the difference between their highest and lowest free energy at a minimum conformation was higher than 10 kcal/mol, calculated with B3LYP/aug-cc-pVTZ. Names are in accordance with Figure **??**.

**Appendix 0 Figure S.I.5.**
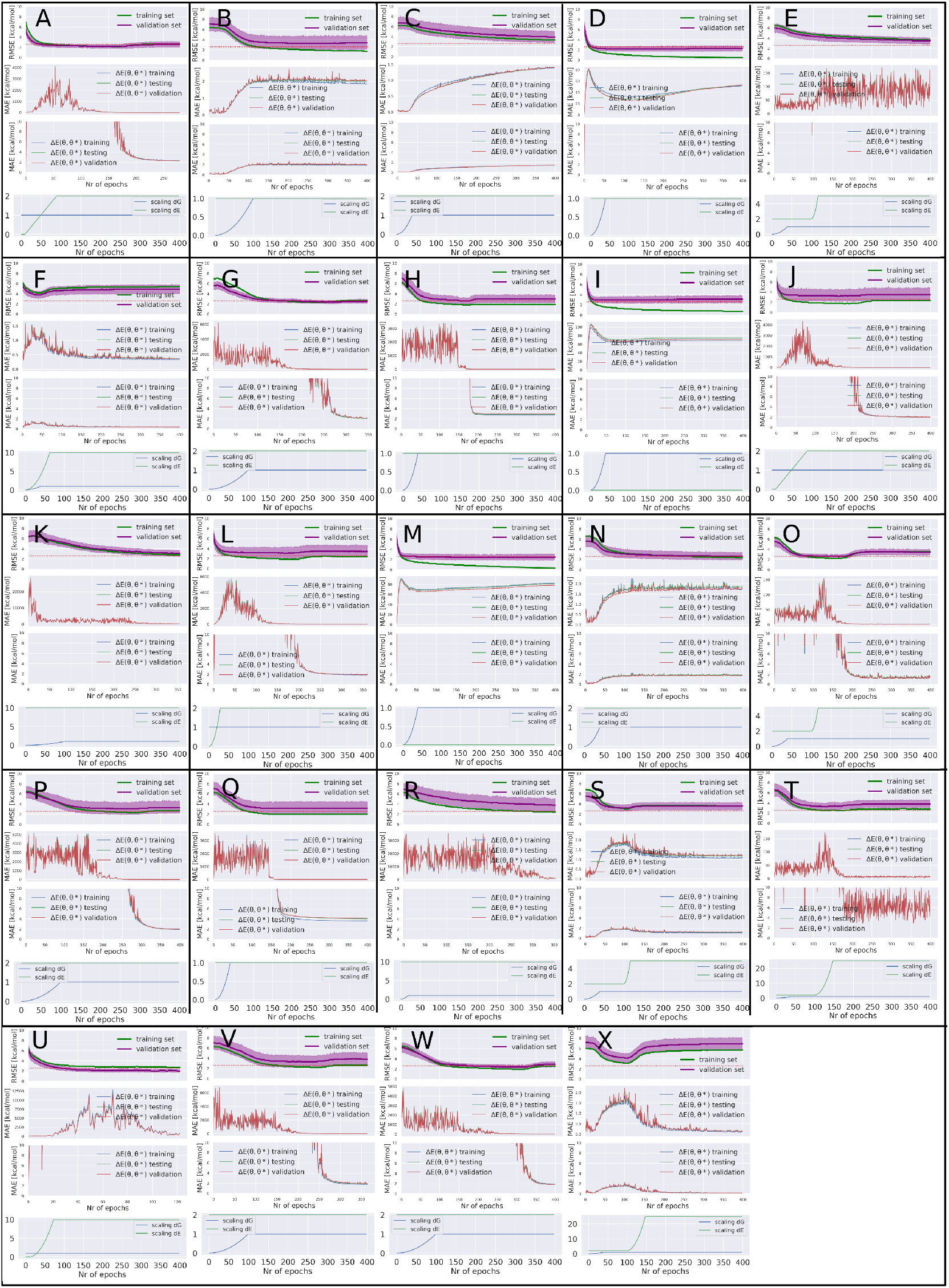
The top panel of the figure shows the RMSE of the training/validation set. The red dotted line indicates the validation set performance of final results shown in Figure 8. The two middle panels show the Δ*E*(*θ, θ*^*^) for the coordinate sets in the training (blue), validation (red) and test set (green). The top middle panel shows the full range of the values on the y-axis, while the bottom middle panel is limited to the interval [0,10] kcal/mol. The bottom panel shows the scaling variables used to scale the two terms (free energy and energy deviation) of the molecular loss function. In addition to the scaling variables used in the loss function there were three hyper parameters controlling the performance of the optimizer: the learning rate (LR) for the SGD and AdamW optimizer (optimizing the bias and the weight of the neural net) and the weight decay. The following list contains the LR for the training runs shown, if weight decay or a learning rate reduction method was used it is explicitly mentioned. **A, H, J, K, L, M, R, U, V, W**: AdamW: LR of 1e-4, SGD: LR of 1e-4. **B** : LR AdamW: 1e-4, LR SGD: 1e-9. **F, I, N, X, S**: LR AdamW: 1e-4, LR SGD: 0. **D**: LR AdamW 1e-5, LR SGD: 0. **O**: LR AdamW 1e-4, LR SGD: 1e-6. **E** : LR AdamW 1e-5, LR SGD: 1e-6. **G** : LR AdamW 1e-4, LR SGD: 1e-4, weight decay: 1e-05. **Q** : LR AdamW 1e-4, LR SGD: 1e-4, with LRReduction on Plateau for AdamW. **T** : LR AdamW 1e-4, LR SGD: 1e-6, weight decay: 1e-9.

**Appendix 0 Figure S.I.6.**
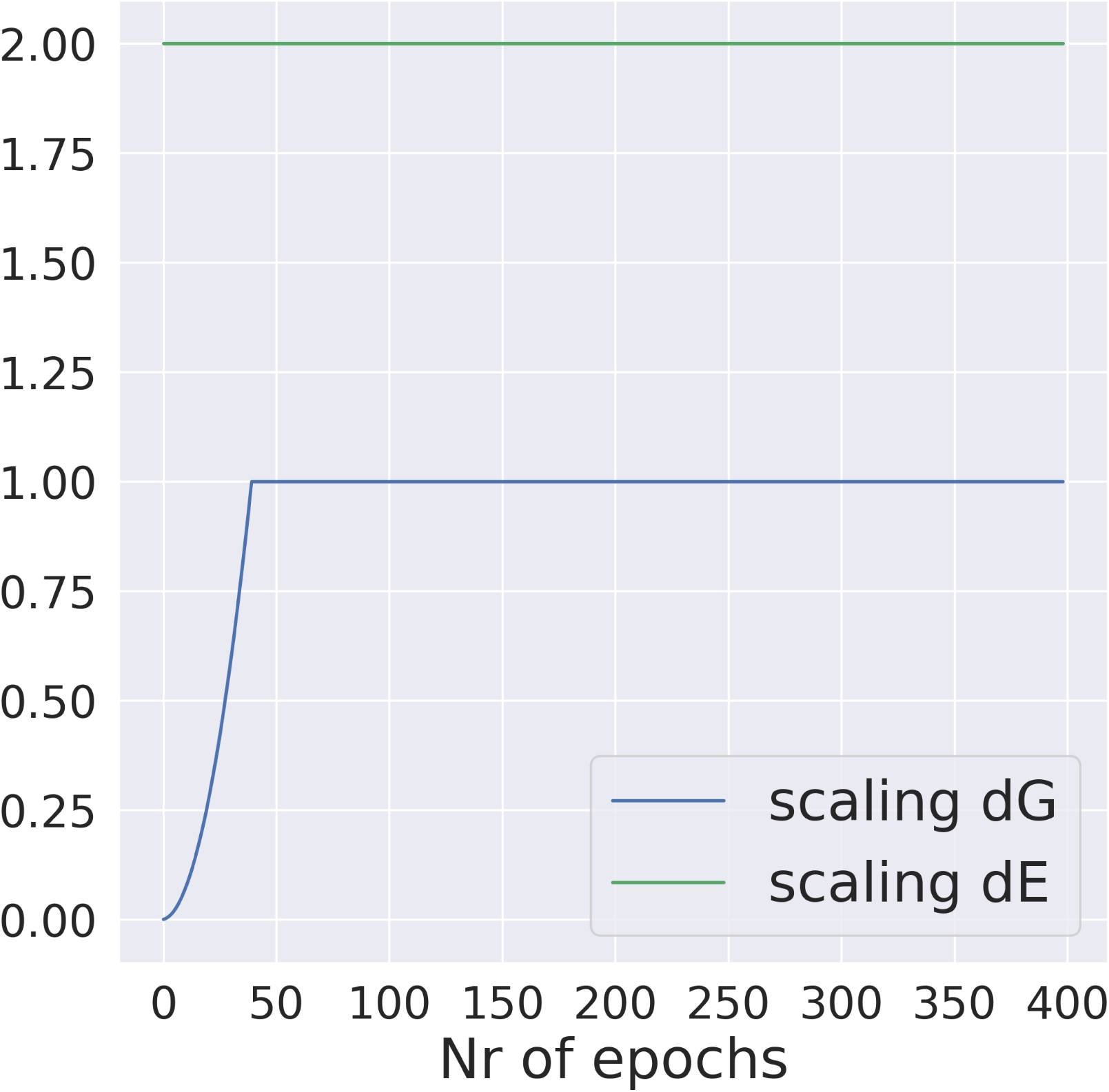
The scaling factors for f(epoch) and g(epoch) used in the molecular loss function in the reported results in 8.

**Appendix 0 Figure S.I.7.**
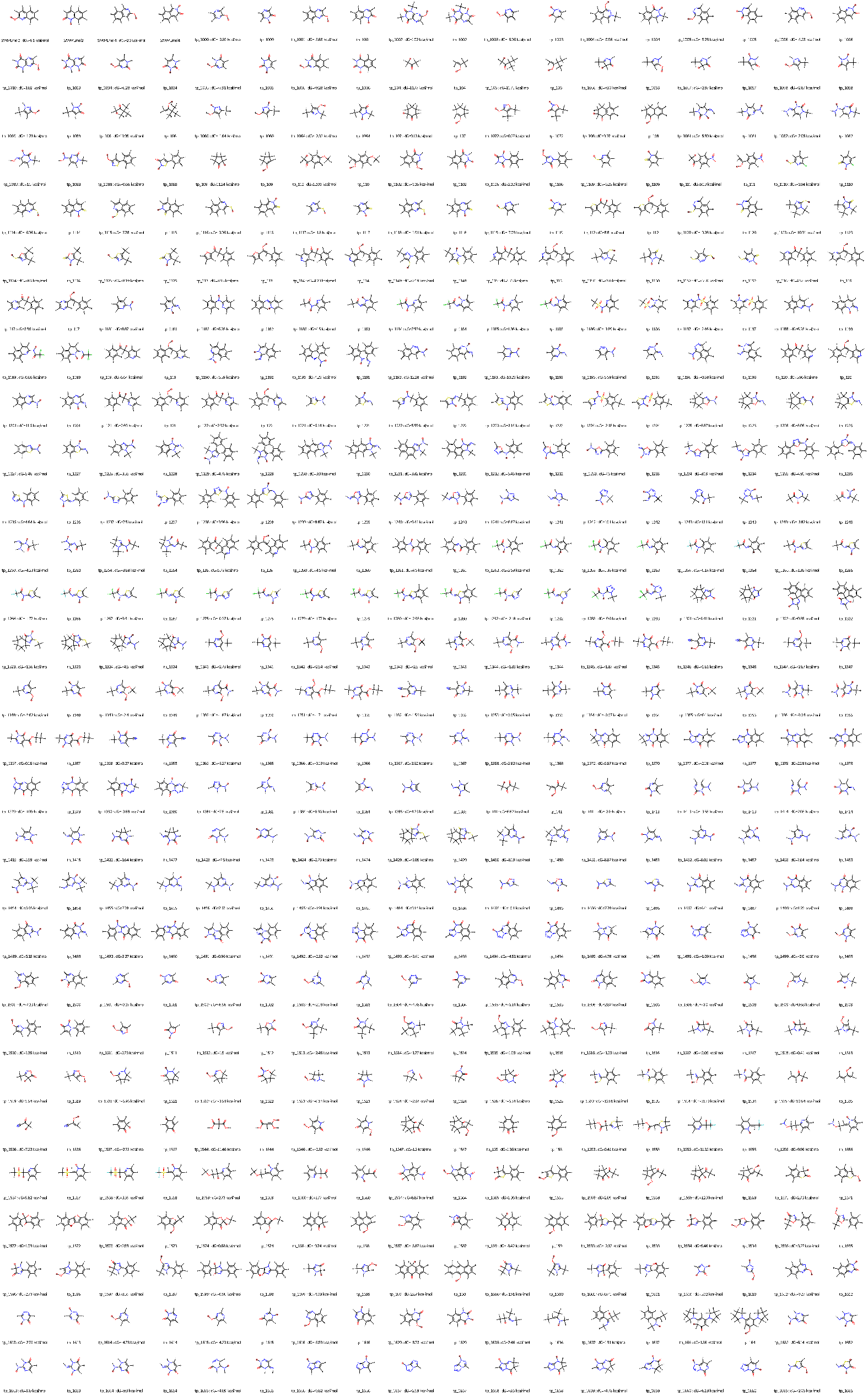
The full tautomer set is shown (part 1). The hydrogen that is moved in the reaction is highlighted in red, *dG* indicates the experimental free energy difference in solution.

**Appendix 0 Figure S.I.8.**
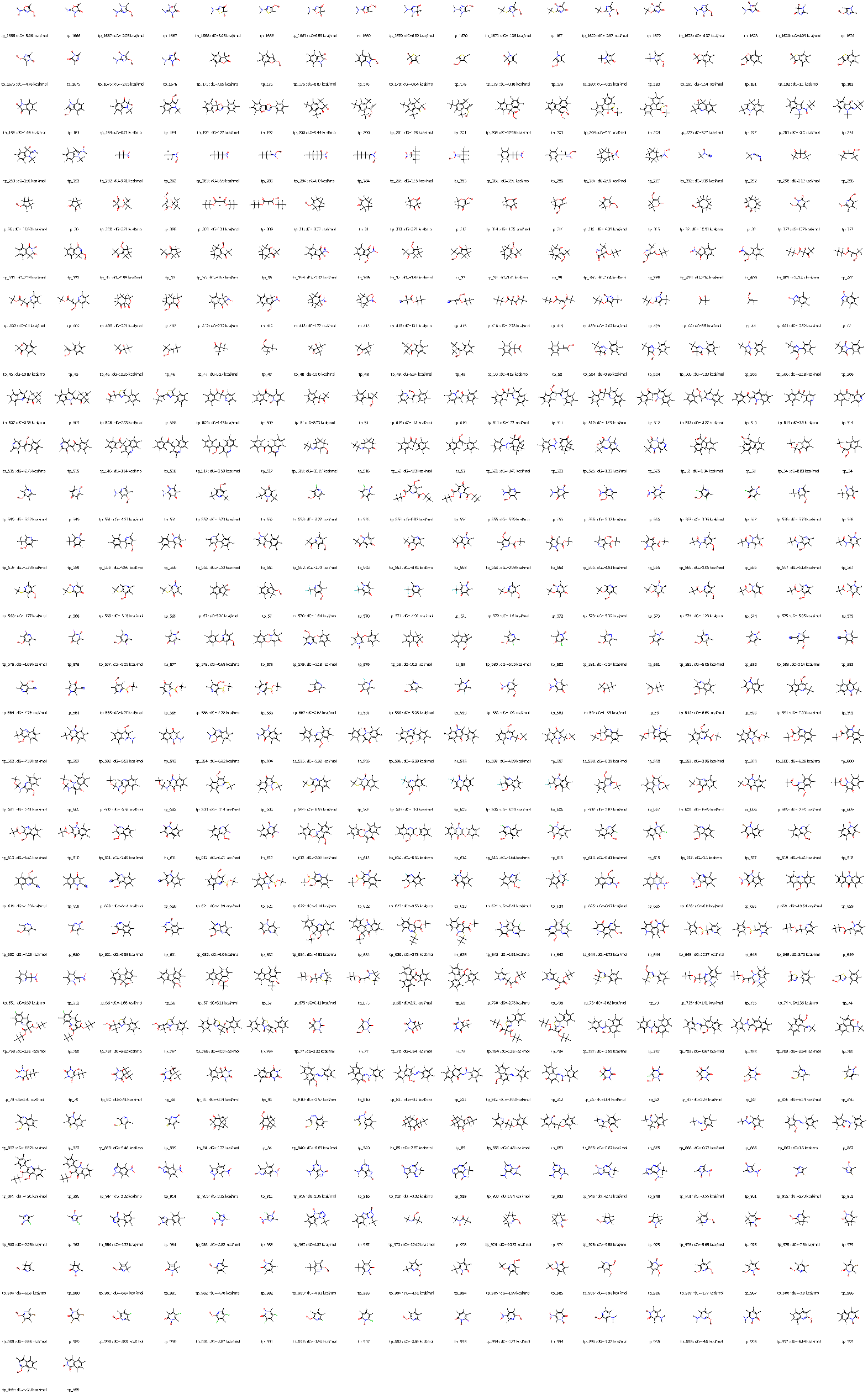
The full tautomer set is shown (part 2). The hydrogen that is moved in the reaction is highlighted in red, *dG* indicates the experimental free energy difference in solution.

## Footnotes

1 (https://qcarchive.molssi.org/apps/ml_datasets/

2 https://aiqm.github.io/torchani/examples/nnp_training.html

